# scBaseCount: an AI agent-curated, uniformly processed, and autonomously updated single cell data repository

**DOI:** 10.1101/2025.02.27.640494

**Authors:** Nicholas D. Youngblut, Christopher Carpenter, Arshia Nayebnazar, Abhinav Adduri, Rohan Shah, Chiara Ricci-Tam, Jaanak Prashar, Rajesh Ilango, Noam Teyssier, Silvana Konermann, Patrick D. Hsu, Alexander Dobin, Dave P. Burke, Hani Goodarzi, Yusuf H. Roohani

**Affiliations:** Arc Institute; University of California, Berkeley; University of Pennsylvania; Stanford University; University of California, San Francisco

## Abstract

Single-cell RNA sequencing has transformed cell biology by enabling precise transcriptomic measurements of individual cells. The Sequence Read Archive (SRA) is the largest public repository of sequencing reads, yet much of it remains underutilized due to unstandardized metadata and the cost of processing reads. Here, we introduce scBaseCount, a single-cell RNA sequencing database that leverages an AI agent to automate discovery and metadata extraction, and standardize data processing. Built by directly mining all 10x Genomics datasets from SRA, scBaseCount is the largest freely accessible public repository of single-cell gene expression data, comprising over 502 million cells across 27 organisms and 75 tissues, offering an unbiased view of the composition of data within SRA. Uniform processing enables measurement of both intronic and exonic reads, non-coding gene expression and improves alignment across experiments as well as the performance of AI models trained on this phenotypically diverse data. Moreover, scBaseCount provides a blueprint for how AI can be leveraged to curate and autonomously update large biological data repositories.

## 1. Introduction

The ability to precisely measure the transcriptomic state of individual cells has transformed the study of cell biology. By uncovering the heterogeneous states of cells as they engage in diverse processes and functions across species and tissue contexts, single-cell RNA sequencing has revealed the finer details of cell identity and behavior that were previously inaccessible through bulk approaches. These discoveries have significantly influenced various fields, from developmental biology to cancer research (Rood et al., 2022; Eraslan et al., 2022; Melms et al., 2021).

Over the past decade, interest in integrating single-cell datasets across institutions and research laboratories has grown significantly. Notable efforts, such as the Human Cell Atlas (Rozenblatt-Rosen et al., 2017) and the CZ CELLxGENE dataset (CZI Cell Science Program et al., 2025), have substantially expanded the availability of curated single-cell RNA sequencing (scRNA-seq) datasets. These efforts have proven instrumental in advancing our understanding of cell identity, cellular development and differentiation, and disease mechanisms, while also providing valuable training data for AI-driven modeling of cellular states. These models are designed to capture context-dependent cellular function and behavior and predict cell state in experimentally unobserved settings such as in response to perturbation (Lopez et al., 2018; Theodoris et al., 2023; Cui et al., 2024; Hao et al., 2024; Rosen et al., 2023; Pearce et al., 2025; Adduri et al., 2025). Building these models has emerged as a major frontier in the application of AI to biology (Bunne et al., 2024; Roohani et al., 2025).

However, initiatives like CELLxGENE and the Human Cell Atlas primarily rely on contributed datasets, which is only a subset of the publicly available data that is accessible through the NIH-hosted Sequence Read Archive (SRA)—the largest repository of raw single-cell sequencing data (Leinonen et al., 2011). While this current approach allows for deeper, expert-level dataset curation and annotation, it also limits the scale and diversity of data available for analysis, particularly for AI models, which typically do not rely on cell labeling. This underscores the need for a new approach to single-cell genomics data curation, one that operates at the scale required for AI model training and is free from the constraints of manual dataset annotation.

Another challenge with existing repositories of single cell data is that aggregation of datasets from diverse sources introduces analytical variability due to differences in alignment tools, reference genomes, and read counting strategies. In the bulk RNA-seq domain, large-scale reanalysis efforts like the ReCount initiative have previously demonstrated the power of standardized pipelines in minimizing these analytical confounders (Frazee et al., 2011). By uniformly reprocessing bulk RNA sequencing data, ReCount provided researchers with a resource where biological variation was not impacted by inconsistencies in data processing pipelines. Drawing on these lessons, we recognized the need for a similar data repository for single-cell genomics, spanning a wide range of species and tissues while adhering to a consistent processing standard. Such a resource should enable more robust cross-study comparisons, facilitate meta-analyses, and better support AI-driven research aimed at modeling cell behavior across diverse biological contexts.

Here, we present scBaseCount, a single-cell genomics data repository that represents the largest-scale reprocessing effort of scRNA-seq data to date. By directly mining all publicly available 10x Genomics datasets from SRA, scBaseCount enables unbiased and comprehensive single-cell transcriptomic analysis of processed data from more than 502 million cells spanning 27 organisms and 75 tissues. Uniform processing enables reduction of processing artifacts and improved analysis downstream, while also producing the largest resource of non-coding transcriptomic data at the single cell level. When used to train a cell embedding model, scBaseCount enables meaningful improvements in discriminating between diverse phenotypes. scBaseCount is curated by an AI agent and is autonomously updated as more data becomes available in the repository. This approach opens many new avenues for AI-powered curation and maintenance of large-scale data repositories, offering a generalizable solution to challenges of metadata standardization that currently limit access to many biological datasets.

## 2. Results

### 2.1. scBaseCount, the largest public repository of 10x single-cell data provides novels insights on SRA data composition

scBaseCount is the first comprehensive single cell database built by directly mining all publicly accessible 10x Genomics scRNA-seq data from the Sequence Read Archive (SRA) and applying a standardized processing pipeline to improve data harmonization. Using an AI-driven agent, `SRAgent`, we systematically and continuously identify repositories, unify metadata across diverse sources, and facilitate the discovery, annotation, and reprocessing of raw single-cell RNA-seq data.

To date, our `SRAgent` has processed 208,939 SRA experiments (i.e. SRX entries), of which 105,343 were identified as 10x Genomics sequencing libraries. With the inclusion of 6,059 additional samples currently part of CZ CELLxGENE (CZI Cell Science Program et al., 2025), we have identified a total of 111,402 target SRX entries at the time of this writing. So far, we have reprocessed 61,381 entries.

Currently, scBaseCount comprises over 502 million cells, with an average of 6052 unique molecular identifiers (UMIs) per cell. When compared to other large repositories of single cell data, such as CZ CELLxGENE (123 million cells) and Human Cell Atlas (65 million cells), scBaseCount is the largest collection of publicly available single-cell datasets at the time of this writing (**Figure 1A**). With data from 27 organisms and 75 tissues, scBaseCount provides a significantly broader range of experimental contexts compared to the current largest repository, CZ CELLxGENE (**Figure 1B,C**).

**Figure 1.**
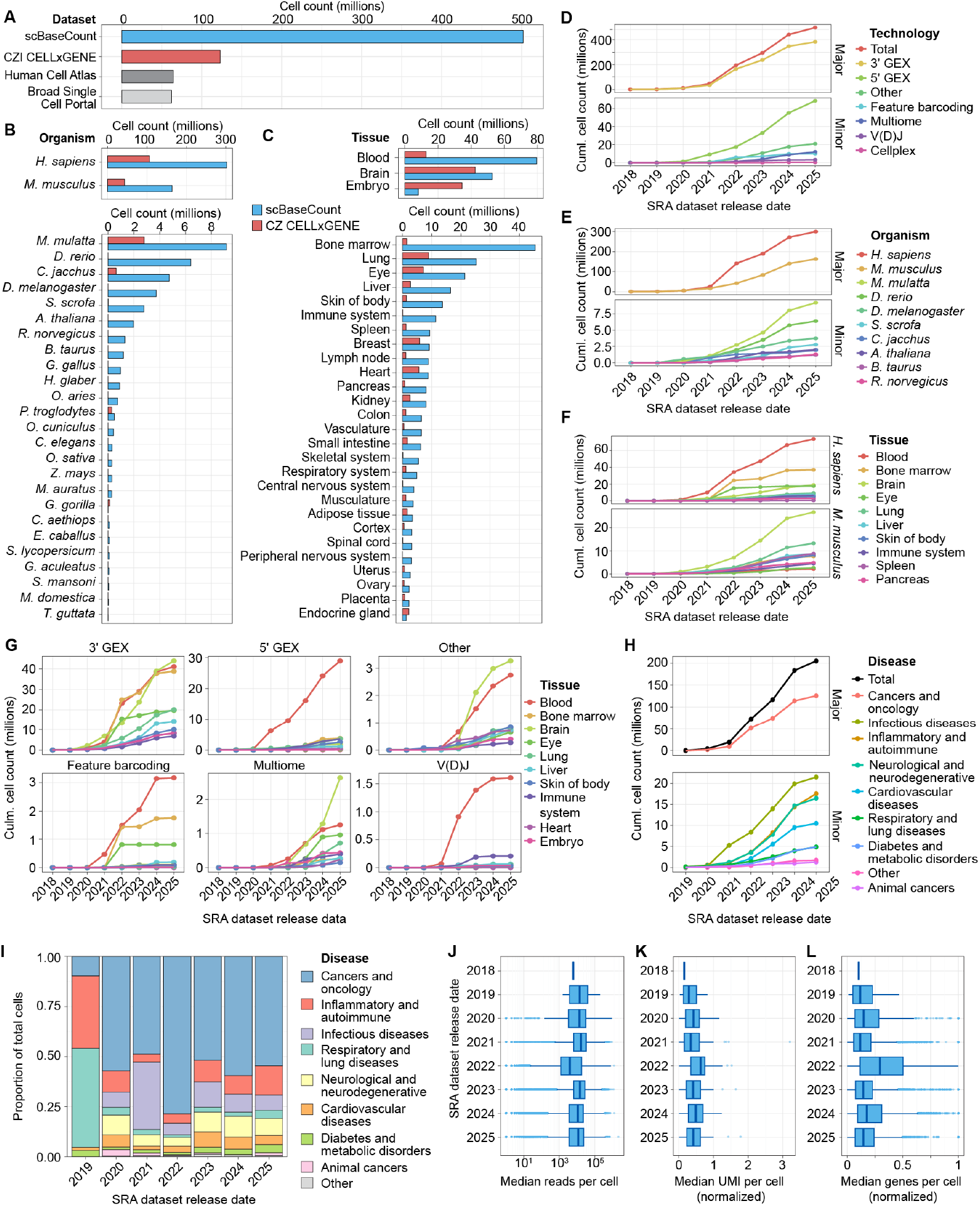
scBaseCount: The largest publicly available repository for single-cell gene expression datasets across species and tissues. (**A**) Comparison of total cell counts among various cell repositories. (**B**) Comparison of cell counts across species in scBaseCount versus CZ CELLxGENE, highlighting a broader species representation and substantially more cells per organism in scBaseCount. (**C**) Comparison of cell counts across the top 30 tissues (all mammals) in scBaseCount and CZ CELLxGENE (human and mouse), illustrating the increased tissue diversity and representation in scBaseCount. Cumulative cell counts by SRA release date grouped by 10x Genomics chemistry (**D**), organism (**E**), tissue (**F**), tissue and 10x Genomics technology (**G**), or disease (**H**). Only humans and mice are included in **H**. For the sake of clarity, only the top 8 to 10 categories with the most total cells are shown if > 10 exist. (**I**) The proportion of cells by disease category by year from 2019 to 2025.(**J**) The distribution of reads per cell. (**K, L**) The distribution of UMIs or genes per cell, normalized to reads per cell.

In addition to dataset identification, `SRAgent` also attempts to extract key metadata for each SRX, including 10x chemistry, cell vs nuclei suspension type, and associated diseases and tissues. Given the widespread use of SRA as a repository for single-cell RNA sequencing data, a comprehensive curation of its contents and associated metadata offers rare insight into broader trends in the field, supported by concrete numbers. We observe that 3’ data has dominated, accounting for 77% of all cells in our collection (**Figure 1D**). This dominance has remained consistent over the past seven years, although 5’ data has shown strong growth in the last two years (13% of cells in 2024-2025). Among the organisms in scBaseCount, human and mouse cells account for 60% and 32%, respectively (**Figure 1E**). Notably, mouse cells have increased proportionally 24% from 2023 to 2025, while human cells declined by 29%. Among other organisms, recent trends over the past three years indicate increasing representation of Macaca mulatta (macaque) and Danio rerio (zebrafish). Among tissues, blood and bone marrow are the largest share of the data, while brain data has shown strong growth within the past few years (**Figure 1F**). Notably, some tissues are strongly contingent on 10x chemistry, such as blood dominating the 5’ and V(D)J data, which makes sense given their relevance to immune profiling (**Figure 1G**). Among human and mouse datasets, the “Cancers and Oncology” disease category dominated, and comprises 30.7% of all cells (**Figure 1H**). The proportion of cells per disease category has shifted dramatically in some cases between years, such as infection disease spiking to 33.7% in 2021 from 7.7% in 2020 (a time period overlapping with the peak of the COVID-19 pandemic) and then declining to 5.9% in 2022 (**Figure 1I**). These data highlight how trends in biomedical research have shifted.

While the number of cells has dramatically increased over the past few years, the sequencing depth per cell has been relatively constant (**Figure 1J**). We found no substantial increase in the number of genes or UMIs per cell after normalizing by sequencing depth (**Figure 1K,L**). The findings suggest that while sequencing costs have declined and sequencer throughput has increased, researchers have opted for sequencing more cells versus increasing the depth per cell.

### 2.2. Automated single-cell data discovery and annotation using SRAgent

To systematically identify and integrate single-cell RNA-sequencing datasets, we developed `SRAgent`, a hierarchical agentic workflow built with LangGraph around large language models (LLMs) and specialized tools for querying the SRA. Specifically, `SRAgent` employs a hierarchical workflow of ReAct agents (Yao et al., 2022) that asynchronously access `eSearch`, `eSummary`, `eFetch`, `eLink`, NCBI HTML scraping, SRA BigQuery, `sra-stat`, and `fastq-dump`. This workflow continuously mines publicly accessible SRA datasets, retrieves key metadata (e.g., organism, tissue, and disease), and stores these annotations in a relational database (**Figure 2A**). This automated approach enables rapid discovery of new studies while ensuring metadata curation remains consistent and scalable.

**Figure 2.**
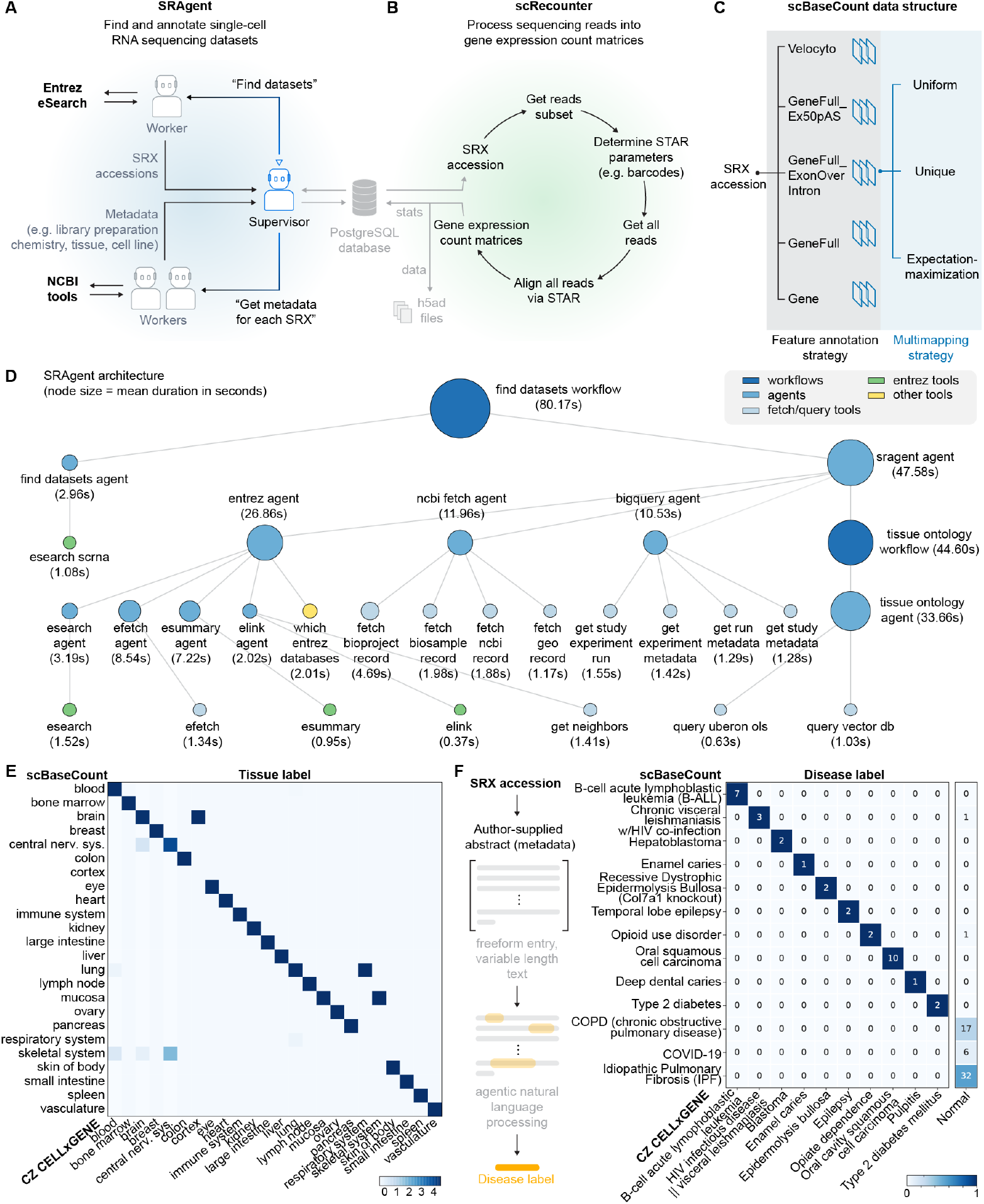
AI-driven curation and standardized processing of single-cell RNA-seq data in scBaseCount. (**A**) SRAgent workflow: A hierarchical AI-driven pipeline for automated dataset discovery and metadata curation from the SRA. SRAgent systematically queries NCBI tools (e.g., eSearch, eFetch) to identify 10x Genomics datasets, retrieve metadata (e.g., tissue type, library preparation chemistry), and store structured annotations in a SQL database. (**B**) scRecounter pipeline: A Nextflow-based workflow for processing raw single-cell sequencing reads into gene expression count matrices. scRecounter downloads and aligns sequencing reads using STARsolo, automatically detects optimal barcode parameters, and generates harmonized expression matrices stored in h5ad format. Process tracking is managed via a PostgreSQL database hosted on GCP. *(caption continued on next page)* (**C**) `scRecounter` uses multiple feature annotation and multi-mapping strategies to generate a variety of CELLxGENE count tables. Users can choose the option that best suits their application. We currently provide access to all variations in scBaseCount. (**D**) Simplified diagram of the hierarchical agentic architecture of SRAgent. Node size denotes the average of time required per workflow component. (**E**) Matching of SRA Agent tissue labels with CELLxGENE tissue labels for a shared subset of data. (**F**) Matching of SRX disease labels derived from author-supplied abstracts at the dataset level, with CELLxGENE disease labels, which are generally assigned to subsets of cells.

Our simplified diagram of the agent-tool hierarchy illustrates the sophistication of our framework (**Figure 2B**). While LLMs are rapidly increasing in agentic abilities (METR: https://metr.org/blog/2025-03-19-measuring-ai-ability-to-complete-long-tasks/), we found that the overall task for `SRAgent` and the number of tools required were too complicated for a single ReAct agent. We instead decomposed the work into sub-tasks handled by a supervisor-worker agentic architecture. For instance, the `find-datasets` agent will first call `eSearch` to obtain new datasets and then the Supervisor calls the `SRAgent` agent to obtain the target metadata. The `SRAgent` agent can call the the `entrez`, `ncbi_fetch`, `bigquery`, and `tissue_ontology` agents. Moreover, the `entrez` agent can call a set of sub-agents that each utilize specific Entrez tools. This architecture reduces the task complexity for any given agent and is more token efficient, since only the final result of an agent is passed back to the supervisor.

In total, `SRAgent` has processed 208,939 datasets, of which 105,343 were identified as 10x Genomics sequencing libraries. Assuming a human could process 1 dataset per minute, 3482 hours (145 days) would be required to analyze all 208,939 datasets. We estimate a total token usage of 13.2 × 10^12^ at a cost of $17,000. By using `SRAgent` to identify single cell datasets, and then passing them onto `scRecounter`, we estimate that greater than 0.5M CPU hours were saved compared to if we had attempted to brute force the task by running every SRX accession through the 10x barcode detector in `scRecounter`.

When comparing `SRAgent`’s automated tissue annotations to those in CZ CELLxGENE, we found that in most cases, the agent accurately extracted the correct tissue labels (**Figure 2E**, Table S4). This observation aligns with recent studies demonstrating the effectiveness of large language models in cell type annotation (Kazmi et al., 2025; Hou and Ji, 2024; Chen and Zou, 2024; Xiao et al., 2024). While AI models of cell state do not rely on these labels during training, `SRAgent`’s ability to perform reliable tissue labeling enhances confidence in its data curation and annotation capabilities.

In addition to tissue annotations, `SRAgent` assigns a disease label to each SRX based on the abstract provided by the authors when submitting their data to SRA (Fig.2F). To validate these disease labels, we compared them to those in CZ CELLxGENE for the subset of datasets shared between the two repositories. Because SRAgent labels are derived solely from the uploaded abstract, they reflect the condition under study by the data generator, whereas CZ CELLxGENE typically indicates whether the individual cells themselves are diseased. Thus, comparing diseased-cell annotations in CZ CELLxGENE against SRAgent labels provides a reasonable validation, since diseased cells are likely to appear in datasets studying that disease. SRAgent successfully identified the studied disease for all such datasets (Fig.2F), S4. Beyond this, even in datasets where cells are labeled as “Normal” in CZ CELLxGENE, SRAgent sometimes detected that the study may nevertheless focus on a disease. We manually examined several such cases and found strong evidence supporting the assigned label S4, S3, S2. These datasets are particularly valuable for researchers investigating rare or understudied conditions, which have not yet been profiled at scale in large atlases but can be reconstructed from smaller datasets distributed across the SRA.

As part of this publication, we have made the `SRAgent` code publicly available. The research community can access and utilize `SRAgent` to further expand single-cell dataset discovery and integration efforts. Of note, `SRAgent` is a general tool that can provide information for many SRA-related queries and includes an assortment of CLI tools for ease-of-use.

### 2.3. Standardized data re-processing using scRecounter

To process the 10x Genomics datasets identified by `SRAgent`, we developed `scRecounter`, a custom `Nextflow` pipeline (Di Tommaso et al., 2017) designed for efficient and scalable single-cell data reprocessing. To date, the pipeline has mapped a total of 13.8 × 10^12^ reads with `STARsolo`.

In analyzing scRNA-seq data, researchers often make choices regarding gene models and counting strategies, as there is no universal approach that fits all applications. Depending on the specific analysis, whether focused on standard gene expression quantification, alternative splicing, or transcriptional dynamics, different gene models and counting strategies may be more appropriate. With `scRecounter`, our goal is to provide a comprehensive set of possibilities, enabling researchers to select the most suitable approach for their own study. To achieve this, `scRecounter` offers multiple feature annotation strategies, including Gene, GeneFull, GeneFull_Ex50pAS, GeneFull_ExonOverIntron, and Velocyto (La Manno et al., 2018b) (see Methods). These models vary in their treatment of exonic and intronic regions, allowing users to choose between exon only counts (Gene), exon and intronic counts (GeneFull), the hybrid between the two that is implemented in CellRanger, (GeneFull_Ex50pAS), or a combination of the two for RNA velocity analysis (Velocyto) (**Figure 2C**).

Additionally, `scRecounter` is flexible in terms of multi-mapping strategies, which determine how multimapped reads are handled. These include expectation-maximization (EM) (Li et al., 2009), which probabilistically assigns multi-mapped reads, as well as unique and uniform strategies, which either retain only uniquely mapped reads or distribute multi-mapped reads evenly (**Figure 2C**).

By incorporating these diverse options, `scRecounter` ensures that scBaseCount remains a versatile resource, accommodating a broad range of single-cell RNA-seq applications while minimizing technical biases. As part of this publication, we have also made the `scRecounter` code available.

### 2.4. Clustering and Annotation of scBaseCount data

Unlike existing single-cell data repositories and atlases, cell annotations in scBaseCount were not manually assigned by independent experts. This presents unique challenges in interpreting the data, understanding its diversity, and exploring its finer structure beyond what is provided in the original SRA annotations. A deeper analysis of the dataset requires comprehensive annotation of cell categories across its entirety. To address this, we developed a computational approach to annotate all data contained within the repository.

Given the diversity of species in the scBaseCount data, we required a cell type annotation strategy that would work consistently across species. For this reason, we embedded the cells using State (Adduri et al., 2025), a state-of-the-art cell embedding model that leverages ESM2 protein embeddings to generate species-agnostic gene representations and has been shown to outperform existing models in differentiating between subtle phenotypes. In order to annotate cell types, embeddings generated for human cells using State were annotated using a cell type classifier trained on data from Tabula Sapiens (Consortium^*^ et al., 2022). Our classifier strategy assigned each cell in scBaseCount to the best matching cell type in Tabula Sapiens within the same tissue, using the tissue labels provided by `SRAgent`. By utilizing both the tissue information, as well as the transcriptional profile, this strategy helps to eliminate artifactual cell type categorizations.

When visualizing the distribution of classified cells across a random sampling of SRXs from scBaseCount, we observed clustering of shared cell types, such as leukocytes and muscle cells, across diverse tissues (**Figure 3A**). Across the full dataset, leukocytes, particularly T-cells and B-cells, were overrepresented in nearly all tissues (**Figure 3B**). This enrichment may reflect sampling biases across studies or a preponderance of diseased samples enriched for immune cell populations. Other commonly observed cell types include epithelial and endothelial cells.

**Figure 3.**
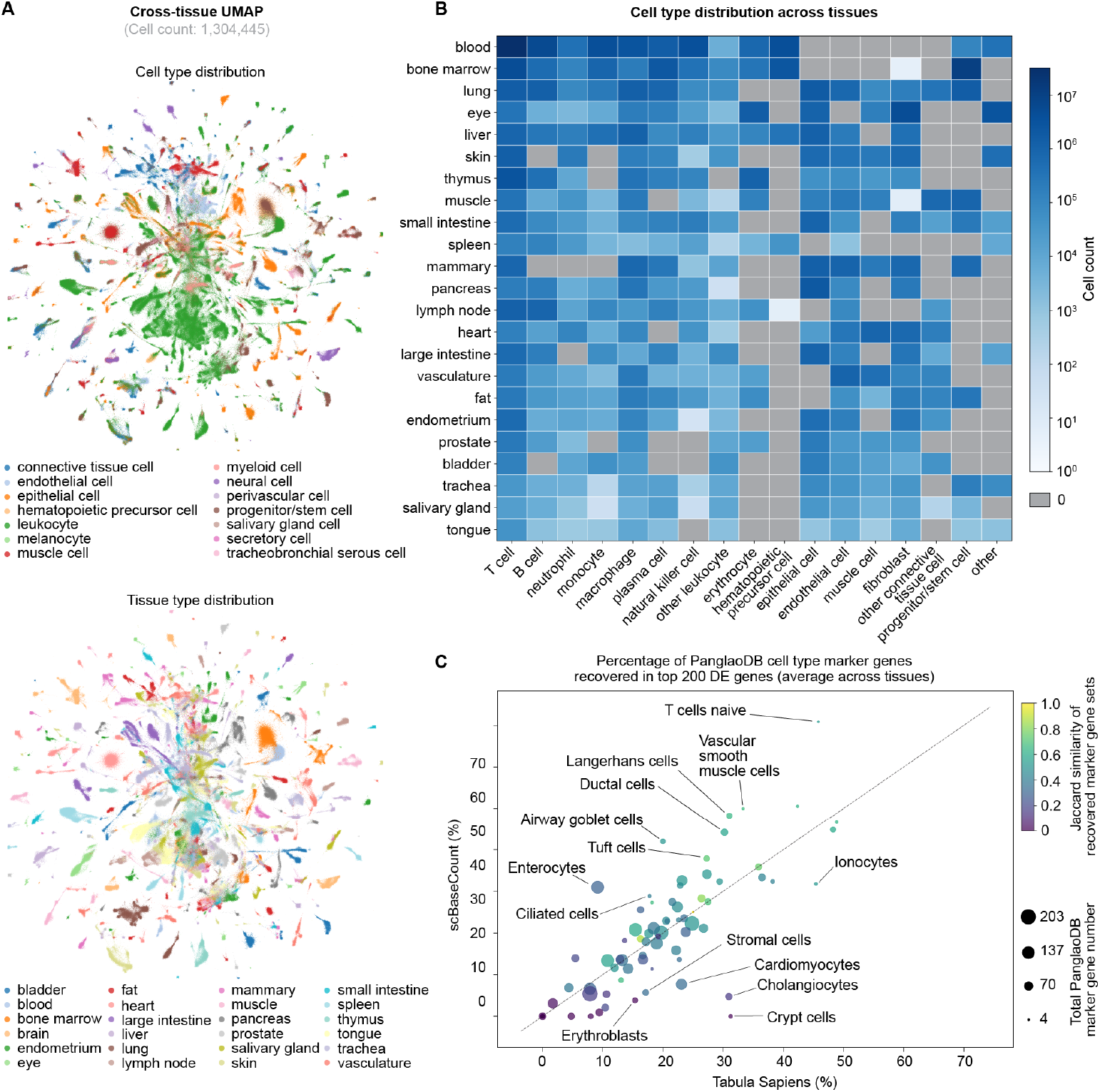
Cell type annotation and clustering of data in scBaseCount. (**A**) UMAP visualization of random per-tissue samples of approximately 50,000 annotated human cells each in scBaseCount (N = 1,304,445) using State embeddings. Outlier cells beyond the 0.1–99.9 percentile range for both x- and y-axis coordinates are excluded from the plot. Top panel: cells colored by predicted cell type; bottom panel: cells colored by tissue label. (**B**) Distribution of inferred cell types by tissue category. (**C**) Scatter plot comparing the percentage of PanglaoDB marker genes recovered among the top 200 upregulated genes for each cell type category between scBaseCount and Tabula Sapiens, averaged across tissues. Differentially expressed genes were computed on random samples of approximately 500,000 annotated human cells for every tissue.

To validate our cell type assignments, we assessed whether known marker genes could be re-captured from the data. Using a t-test differential expression framework applied to random tissue-specific subsets of scBaseCount, we recovered a high proportion of independently curated marker genes from PangaloDB (Franzén et al., 2019). Moreover, when comparing marker gene recovery between Tabula Sapiens and scBaseCount, performance was competitive, even though the classifier itself did not use marker genes, and the curated marker set was independent of the Tabula Sapiens annotations (**Figure 3C**). The overlap between known cell type markers and those recovered from scBaseCount validates both our classifier and the dataset itself. In particular, the shared marker gene signatures between scBaseCount and PanglaoDB indicate that the underlying data in scBaseCount is of sufficient experimental quality to recover biologically meaningful genes, and that it is competitive with Tabula Sapiens as a representation of human cell biology.

### 2.5. Uniform data re-processing reduces the contribution of analytical confounders

A key advantage of a uniform processing pipeline for single-cell transcriptomic data is its ability to minimize technical variability introduced by differences in analytical pipelines. To assess the amount of signal in our data originating from technical factors, we used silhouette scoring (Büttner et al., 2019), a metric that quantifies how well individual data points cluster based on a given categorical variable. The silhouette score ranges from −1 to 1, where higher values indicate that cells are more cohesively grouped within the same category and well-separated from other categories, while lower values suggest overlapping or poorly defined clusters. We applied this approach to evaluate how various categorical metadata variables influence gene expression patterns in scBaseCount. Some factors primarily reflect biological variation (e.g., tissue type), while others capture a mix of technical and biological variation (e.g., sample suspension type, library preparation chemistry, or sample ID). Our goal was to quantify the extent to which groupings based on each factor shape the dataset’s structure and contribute to observed variation in gene expression. In scBaseCount, technical factors such as library preparation chemistry and sample suspension type (single-cell vs. single-nucleus sequencing) exhibited comparable or lower silhouette scores than biologically meaningful categories like tissue type (**Figure 4A**) when compared to CZ CELLxGENE which is composed of pre-processed datasets that are not uniformly processed. For example, in the case of sample ID (largely equivalent to the SRA study or SRP ID in scBaseCount; **Figure S5**) the silhouette scores are similar to tissue type in scBaseCount, while in CZ CELLxGENE, they are significantly higher, suggesting that study-specific processing introduces stronger batch effects (**Figure 4A**).

**Figure 4.**
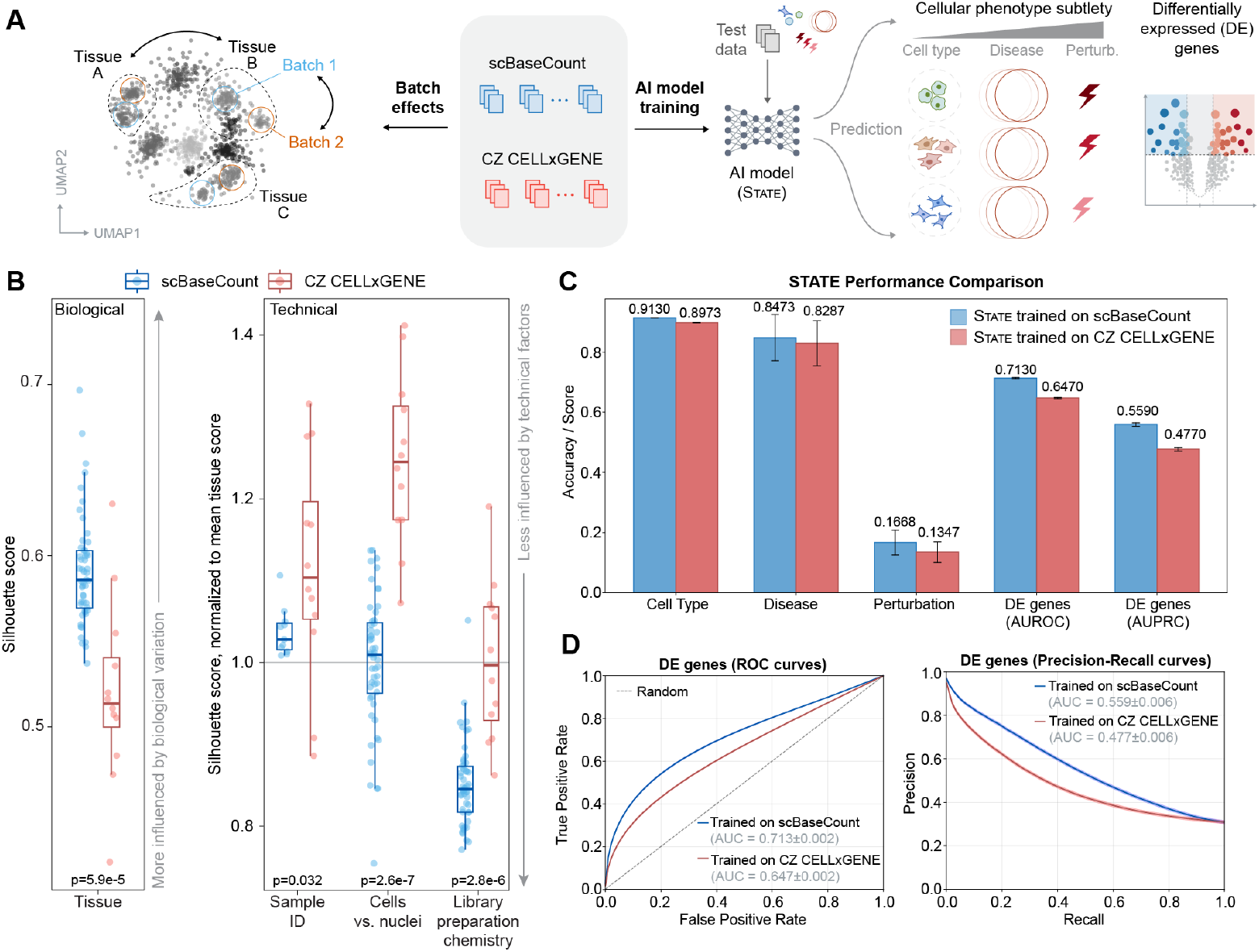
Impact of scBaseCount on downstream analyses and AI model training. (**A**) Comparing the impact of technical and biological factors on clustering in scBaseCount and CZ CELLxGENE, and on downstream AI model training. (**B**) Silhouette scores computed across random subsets of 250,000 cells from scBaseCount (blue) and CZ CELLxGENE (red) for different metadata factors. Higher scores indicate a greater influence of the corresponding factor on data clustering. For technical factors, scores are normalized to the mean tissue score for each dataset. CZ CELLxGENE subsets were drawn from 20 individual human datasets and downsampled to 250,000 cells in two steps, and scBaseCount subsets were randomly sampled from human and mouse accessions. The resulting objects were filtered to retain genes present in at least 10 cells and cells with at least 30 detected genes. All datasets were sum-normalized, log-transformed, and processed using PCA before calculating silhouette scores (`scib_metrics package`). These results demonstrate that, given the contribution of tissue type, technical factors such as sample ID, library preparation chemistry, and sample suspension type contribute less to clustering in scBaseCount compared to CZ CELLxGENE, highlighting how scBaseCount’s uniform processing pipeline reduces analytical variability while preserving biological structure. (**C**) Performance difference in probing cell embeddings from a model (State) trained on scBaseCount versus that trained on CZ CELLxGENE. Embeddings are probed with an MLP to classify specific conditions, like cell type, disease, or perturbation group. (**D**) Performance difference in reconstructing DEGs from a model (State) trained on scBaseCount versus that trained on CZ CELLxGENE. True DEGs, determined as those with FDR < 0.05 using a Wilcoxon rank sum test, are compared to DEG predictions from State model reconstructions (Methods 5.10). Curves are generated by varying the threshold for calling significance from model outputs.

This effect is even more pronounced for sample suspension type, which differentiates single-cell from single-nucleus sequencing. We reasoned that this may likely be due to the common practice of counting singlenucleus data using both exonic and intronic reads (i.e. pre-mRNA annotations), while single-cell data typically includes only exonic reads. While neither approach is inherently incorrect, this inconsistency could introduce strong batch effects when integrating both data types. scBaseCount mitigates this issue by retaining and reporting both exonic and intronic read counts, resulting in better integration of single-nucleus and single-cell datasets.

To further examine this possibility, we focused on four shared datasets, listed in Table S2, that contain both single-cell and single nucleus-sequencing data. We performed Principal Component Analysis (PCA) and visualized the datasets in two-dimensional PC space (**Figure S6A**). In the 2-dimensional PC space, tissue type showed a higher silhouette score in scBaseCount than in CZ CELLxGENE, and conversely, sample suspension type (i.e. cells vs nuclei) scored lower in scBaseCount (**Figure S6B**). Examining the source publications of these four integrated datasets, we found that the single nucleus data in this subset was indeed processed with pre-mRNA references, and the single cells were processed with a standard exonic reference, as is common practice (**Table S2**). Correcting this technical artifact in scBaseCount resulted in better separation of tissue types along the first two PCs compared to CZ CELLxGENE (**Figure S6C**).

Beyond harmonization, uniform data processing in scBaseCount, including consistent treatment of exonic and intronic reads across samples, enables additional downstream analyses, such as RNA velocity, to better model transcriptional dynamics and cell state transitions. As mentioned in the previous section, `scRecounter` provides cell×gene count tables for spliced and unspliced transcripts as well using Velocyto (La Manno et al., 2018a).

### 2.6. Impact of scBaseCount on training AI models of cell state

Next, we evaluated the impact of scBaseCount when used as training data for AI models of cell state, specifically for the task of generating an embedding of the cell that captures meaningful biological variation across diverse contexts. We trained a 100M parameter State Embedding model Adduri et al. (2025) on a filtered subset of scBaseCount and CZ CELLxGENE. State Embedding is an encoder-decoder model that uses a transformer to construct a dense representation of a cell from its gene expression values, and an MLP to reconstruct expression from the dense embedding. After training each model on human cells from scBaseCount and CZ CELLxGENE, we obtained two models, State_bsc and State_cxg, which we then used to embed cells from several held out datasets. We trained classifiers to distinguish between phenotypes (Methods 5.10), using only the cell embeddings from State_bsc and State_cxg, at varying levels of granularity (**Figure 4C**).

At the coarsest level, we evaluated embeddings on cell type classification, as this is the most distinguishable phenotype. For this task, we used the Tabula Sapiens dataset CZI Cell Science Program et al. (2025), which includes data from 24 donors spanning 180 cell types. Both State_bsc and State_cxg achieved very high performance (approximately 90%), though the model trained on scBaseCount showed a modest improvement.

We then tested the models on subtler state changes, such as those induced by disease or perturbation. For disease, we used datasets containing cells infected by COVID-19 Ravindra et al. (2021); van der Wijst et al. (2021); Wu et al. (2024). For perturbation, we used CRISPRi gene knockdown experiments for 2024 essential genes Replogle et al. (2022); Nadig et al. (2025) across four cell lines (HepG2, RPE1, Jurkat, K562). In both cases, State_bsc clearly outperformed State_cxg, improving classification accuracy by 2.24% for infection status and 23.9% for perturbation group.

Finally, to probe even finer phenotypic resolution at the level of individual genes, we compared differential gene expression predictions. Performance on this metric is a strong indicator of a model’s ability to capture perturbation effects, a core requirement for virtual models of cell state and behavior. Here, the benefit of training on scBaseCount was most striking. We conducted a Wilcoxon rank sum test across all perturbation conditions to get a ranked set of differentially expressed genes (DEGs) for each perturbation. We obtained ground truth DEGs by filtering for FDR < 0.05. Next, data from the genetic perturbation screens was embedded and the State Decoder was used to reconstruct the expression for all cells. We obtained ranked lists of DEGs from the State model reconstructions by ordering genes according to the difference in the reconstructed expression for perturbed groups compared to control groups. ROC and PR curves were obtained by varying the sensitivity threshold to call DE (Methods 5.10). State_bsc showed increased sensitivity to subtle gene expression changes (10.2 % AUROC improvement and 17.2% AUPRC improvement over State_cxg) (**Figure 4D**).

Overall, these results highlight the advantages of scBaseCount for training AI models of cell state, that can be attributed both to its larger size and greater species diversity, as well as to its cleaner biological signal enabled by uniform processing across datasets and matched gene sets.

## 3. Discussion

In this study, we present scBaseCount, a continuously updated and uniformly processed multi-species singlecell RNA-seq data repository that is built through directly mining all publicly accessible 10x Genomics datasets. Our motivation stems from the need to create a large, high-quality resource for training computational models of cell state and behavior. Existing single-cell databases rely on manually curated datasets, which limits their size and diversity, failing to leverage the vast reserves of publicly available transcriptomic data. Moreover, integrating individual datasets within these repositories remains a major challenge due to variation in how each of these datasets were processed.

By systematically mining, annotating and processing raw single-cell transcriptomic reads directly from SRA, scBaseCount is designed to be a large, harmonized repository of single-cell datasets with minimal analytical confounders. Inspired by the ReCount initiative, which demonstrated the value of uniformly processed bulk RNA-seq data (Frazee et al., 2011), scBaseCount reduces technical artifacts across studies, ensuring that AI models learn biologically meaningful variation. It also far exceeds other single-cell databases in species diversity, reflecting the breadth of phenotypic information it captures. Since it contains non-coding transcriptional data for all cells, it is currently the largest repository of such data at single-cell resolution.

We believe that these benefits are particularly relevant in the context of training AI models of cell state, where dataset size, species diversity, and disease diversity are crucial to model performance. In our experiments, even when restricting to only human data, training models with scBaseCount results in a marked improvement in a model’s ability to distinguish subtle but important differences in transcriptional state, such as perturbation and infection. Notably, this does not capture the additional performance gains likely to result from scBaseCount’s unprecedented species diversity, as demonstrated in prior work (Rosen et al., 2023; Pearce et al., 2025). Beyond serving as a drop-in replacement for existing datasets, scBaseCount also enables new model architectures. For example, its inclusion of intronic counts across all cells allows modeling of RNA transcriptional dynamics, such as RNA velocity (La Manno et al., 2018b; Lange et al., 2022), at an unprecedented scale.

scBaseCount is the first large biological data repository curated by an AI agent. Our automated workflow, powered by the `SRAgent` AI system and a `Nextflow`-based processing pipeline, ensures consistent metadata curation and analytical treatment across all datasets. This agentic workflow has the advantage of being entirely automated, easily scalable and capable of continuously updating as new data becomes available. Furthermore, this workflow is able to utilize the abstracts associated with data uploads to create meaningful annotations, which are helpful for both biological interpretation and data processing.

Despite these advantages, several challenges remain. While the agentic workflow automates much of metadata extraction, some tasks, e.g. cell type annotation, may require human expertise, community-sourced curation, or additional tools for mapping individual cells onto annotated references (Lopez et al., 2018; Lotfollahi et al., 2022; Domínguez Conde et al., 2022; Heimberg et al., 2024). Other cell-level annotation such as the applied perturbation or donor information are also not accessible through SRA-derived metadata and must be incorporated manually.

In the current release, we have focused on libraries created using the 10x Genomics platform, and sequenced on established Illumina sequencers. As alternative library preparation chemistries become more commonplace and sequencing technologies continue to evolve, we must adapt scBaseCount to maintain compatibility with new modalities, including various chemistries, multi-omic measurements, and spatial transcriptomics. Although our curation to date has prioritized publicly accessible 10x datasets, expanding to other platforms and protected datasets is a key goal. This requires manual requests for access, but we are actively engaging with authors and anticipate that remaining gaps with other repositories will gradually close. Looking ahead, we also plan to extend `scRecounter` to support additional single-cell technologies, further broadening scBaseCount’s scope.

Ultimately, this unified and continuously updated resource provides a foundation for AI-driven efforts to model cell behavior across health and disease. By delivering the largest uniformly processed single-cell data repository, scBaseCount lowers barriers for large-scale model training and integrative analysis. While our focus here is on single-cell applications, many other fields could benefit from centralized, uniformly processed repositories, particularly where standardized metadata are lacking. We believe that AI-driven data curation and automated processing present a compelling new framework for unlocking such datasets. Thus, scBaseCount represents not only a step forward for single-cell modeling, but also a blueprint for building large, uniformly processed repositories in other areas of biology.

## 4. Data and Code Availability

### 4.1. Data availability

All `STARsolo` count matrices are located on Google Cloud Storage at gs://arc-ctc-scBaseCount/202 5-02-25. Documentation on accessing the data can be found at https://github.com/ArcInstitute/arc-virtual-cell-atlas.

### 4.2. Code availability

`SRAgent` and `scRecounter`, and `tiledb-soma-loader` are available on GitHub at https://github.com/ArcInstitute/SRAgent and https://github.com/ArcInstitute/scRecounter, https://github.com/ArcInstitute/tiledb-soma-loader, respectively. Code used for data analysis is available at https://github.com/ArcInstitute/scBaseCount_analysis.

## 5. Methods

### 5.1. SRAgent: Automated dataset discovery and metadata extraction

`SRAgent` is a Python package that utilizes LangGraph for constructing the agentic workflows. We used LangSmith for profiling SRAgent runs.

`SRAgent` is deployed on Google Cloud Platform (GCP) Cloud Run with 2 CPUs and 2 GB of memory per job. To comply with NCBI API rate limits, jobs are triggered every 1-5 minutes, processing 3-5 datasets per run with a peak rate of 300 datasets per hour. The mean processing time per dataset is 80 seconds, with sub-agents and tools called asynchronously for increased parallelization. All extracted metadata is stored in a GCP SQL database for downstream processing.

To obtain Uberon tissue ontologies and MONDO disease ontologies for each SRA dataset (Mungall et al., 2012), we first converted the node text in the Uberon ontology graph to a ChromaDB vector database. `SRAgent` performs a semantic search with a free text tissue description derived from data obtained via NCBI Entrez tools. For each returned record, the agent then explores neighboring terms in the ontology hierarchy and queries the Ontology Lookup Service (OLS) when additional validation is needed. We note that many SRA experiment datasets could not be disambiguated to just one tissue ontology term based on the metadata in NCBI. In such cases, we retained multiple values.

Disease categories were determined from the MONDO ontology graph (Vasilevsky et al., 2022). For computational efficiency, the graph was pruned to just ontology nodes found in the scBaseCount dataset, with intermediate pruned nodes replaced with a single edge in order to retain connections among the remaining nodes. We created a weighted adjacency matrix for all remaining nodes, with weights set by the number of corresponding records in scBaseCount. We then applied the scikit-learn (Pedregosa et al., 2011) spectral clustering algorithm to produce 12 clusters. To create the cluster labels in an unbiased manner, we provided the top 10 ontology IDs per cluster by weight to Claude Opus 4 and prompted for concise cluster labels based on the cluster compositions.

### 5.2. scRecounter pipeline overview

`scRecounter` is a `Nextflow` (Di Tommaso et al., 2017) pipeline designed for scalable reprocessing of singlecell RNA-seq data. So far, we have focused on processing 10X Genomics datasets as identified by `SRAgent`.

The pipeline runs on GCP Cloud Run and submits jobs to GCP Batch. To stay within GCP resource quota limits, the pipeline processes a maximum of three datasets per run, with new runs triggered every three minutes. Process tracking is managed via a PostgreSQL database hosted on GCP.

### 5.3. Identification of 10X chemistry

Once `SRAgent` has predicted that an SRX dataset contains 10X Genomics single-cell data, the dataset is processed through the `scRecounter` pipeline. Since `SRAgent` could not consistently determine specific assay chemistry, the first step of `scRecounter` was to determine the chemistry using a `Cell Ranger`-inspired automated detection algorithm. The first 1 million reads from each SRX are processed using `STARsolo`, testing barcode sets corresponding to 10X 5’, 10X 3’ v2, 10X 3’ v3, and 10X Multiome GEX. 10X 3’ v1 was not included. The barcode file yielding the highest percentage of valid cell barcodes was selected. Datasets with fewer than 30% valid barcodes were removed at this stage.

### 5.4. Reference genome preparation

For gene annotations, we used human and mouse reference genomes from the 10X Genomics repository. `STAR` references for other species were generated using the following workflow: for each species, a widely used ENSEMBL genome assembly was selected and downloaded (Frölich et al., 2023). As with human and mouse annotations, we applied a custom script to filter each species’ GTF file, retaining only relevant biotypes (e.g., “protein-coding” and “lncRNA”) based on their clade (e.g., mammals, birds, fungi). This filtering ensures that only genes measurable by single-cell RNA-seq are included and maintains consistency with 10X Genomics annotations. For `STAR` reference generation, we adjusted the `genomeSAindexNbases` parameter according to genome size, following recommendations from the `STAR` developers (Dobin et al., 2013). These annotations and references are available for download through our portal.

### 5.5. Data processing with STARsolo

After identifying the correct 10X chemistry, we then aligned all SRRs (SRA runs) from an SRX with a single `STARsolo` run, using the following parameters:

~~~
--soloType CB_UMI_Simple
--clipAdapterType CellRanger4
--outFilterScoreMin 30
--soloCBmatchWLtype 1MM_multi_Nbase_psuedocounts
--soloCellFilter EmptyDrops_CR
--soloUMIfiltering MultiGeneUMI_CR
--soloUMIdedup 1MM_CR
--soloFeatures Gene GeneFull GeneFull_ExonOverIntron GeneFull_Ex50pAS Velocyto
--soloMultiMappers EM Uniform
--outSAMtype None
--soloBarcodeReadLength 0
~~~

scRecounter leverages `STARsolo`’s `--soloFeatures` parameter, enabling the generation of various count matrices tailored to specific analytical needs:

- **Gene**: Counts reads mapping entirely to exonic regions of annotated transcripts, capturing fully spliced transcripts. This procedure represents traditional gene expression analyses and was the default in earlier `CellRanger` versions.
- **GeneFull**: Counts reads overlapping entire gene locus, i.e. including exonic and intronic regions, which captures both unspliced (primary) and spliced transcripts and also provides a more comprehensive view of gene activity.
- **GeneFull_ExonOverIntron**: Counts reads overlapping exonic and intronic regions, but assigns higher priority to exonic overlaps. This option helps to resolve reads that map to overlapping genes.
- **GeneFull_Ex50pAS**: Similar to the above option, but with more sophisticated priorities scheme, which prioritizes partial and antisense exonic overlap over intronic reads.
- **Velocyto**: Generates separate count matrices for spliced, unspliced, and ambiguous reads, following the rules from (La Manno et al., 2018b). This enables RNA velocity analyses to infer dynamic cellular processes.

For `Gene`, `GeneFull`, `GeneFull_ExonOverIntron`, and `GeneFull_Ex50pAS`, three multi-mapping strategies are used:

- **Unique**: retain only uniquely mapped reads
- **Uniform**: distribute multi-mapped reads evenly
- **Expectation-Maximization (EM)**: probabilistically assign multi-mapped reads

`Velocyto` generates the following count matrices:

- **Spliced**: reads mapping exclusively to exonic regions, representing mature mRNA molecules
- **Unspliced**: reads containing intronic sequences, representing nascent pre-mRNA transcripts
- **Ambiguous**: reads that cannot be definitively classified as (un)spliced due to mapping ambiguity

In total, the pipeline generated 15 count matrices per SRX dataset.

### 5.6. Reprocessing datasets in CZ CELLxGENE

We obtained accession codes corresponding to CZ CELLxGENE collections and re-processed the corresponding SRXs using the method outlined above. Constituent SRXs from each collection were then assembled into one `h5ad` object per collection, using *Unique GeneFull_Ex50pAS* counts.

We then mapped the observation metadata from each CZ CELLxGENE collection to our re-processed collections by matching cell barcodes. Specifically, within each collection, the cell barcodes follow the format `[ATCG]`-`[10x_well_id]`, and our `STARsolo`-processed data follows the same format. Therefore, for each collection, we mapped the `10x_well_id`s in the CZI version of the collection to ours by assigning each identifier to the corresponding CZ CELLxGENE identifier that maximized the number of intersecting barcodes. By using this method, we were able to successfully map metadata from CZ CELLxGENE collections into scBaseCount.

### 5.7. Evaluation of analytical factors in CZ CELLxGENE and scBaseCount

In an effort to evaluate the contribution of suspension type to the data structure, we selected four human datasets shared between CZ CELLxGENE and scBaseCount collections that contained both single-cell and single-nucleus data. We identified the matching cells in CZ CELLxGENE using the combination of `observation & _joinid` and `cell_type` as unique cell identifiers. We excluded cells with fewer than 100 unique genes and genes observed in fewer than 40 cells. The resulting datasets were then sum-normalized and log-transformed. Principal component analysis (PCA) was performed using 50 principal components (PCs), with the top two PCs visualized using `scanpy`. Silhouette scores were computed using the `scib_metrics` package.

For evaluating organism classification accuracy by `SRAgent`, we used SRA BigQuery as ground-truth. Notably, manual assessment of misclassified datasets revealed that many were xenografts, so mis-classification was somewhat due to the constraint in SRA that “organism” must be a single value.

### 5.8. Dataset subsampling

For CZ CELLxGENE, each subsampled data contained samples from 20 projects, where all projects were solely 10X assays and labeled as ‘primary_data’ in CZI metadata. Each dataset was subsampled to 30% of the original before joining. Each sample was subsequently downsampled to 250, 000 cells as necessary. For scBaseCount, we selected GeneFull_Ex50pAs counts from randomly chosen SRX accessions, and concatenated them together until the object reached 250, 000 cells. In both cases, half of the subsets were mouse, and half were human.

### 5.9. Model training

To evaluate the comparative utility of scBaseCount and CELLxGENE for training single-cell foundation models, we trained the State Embedding model (Adduri et al., 2025) using 108 million trainable parameters. The architecture consisted of an eight-layer transformer with an embedding dimension of 1024 for the encoder and a multi-layer perceptron to decode expression. Training was conducted on four GPUs with a batch size of 96 and gradient accumulation of 8, yielding an effective batch size of 3072. All other training details match the primary manuscript.

For the scBaseCount dataset, we applied a filter to retain only human cells, excluding those with fewer than 300 non-zero gene counts or 500 unique molecular identifiers (UMIs), resulting in 214 million cells for model training. The CELLxGENE dataset was not additionally filtered beyond its original curation, resulting in 134 million cells for model training.

Both datasets were standardized to a common gene set comprising 19,790 human protein-coding genes. The State Embedding model utilizes pretrained gene embeddings as input features, an approach that has demonstrated superior performance compared to one-hot encoding schemes (Rosen et al., 2023; Pearce et al., 2025). We generated gene representations using ESM-2-based featurization, computed by averaging per-amino acid embeddings across all protein-coding transcripts within each gene. To remove dataset size as a conflating factor, models were trained for 100,000 optimization steps rather than for a fixed number of epochs. To ensure unbiased evaluation, Tabula Sapiens data were systematically excluded from both training sets.

### 5.10. Probing embeddings for separability

To assess the impact of the training dataset on cell embedding information content, we used State Embedding models, trained on scBaseCount or CELLxGENE, to embed held out datasets. For each dataset, we trained a simple MLP probe to classify different conditions given only the cell embedding, such as the applied perturbation, the cell type, or the disease state. The MLPs contained two linear layers with one ReLU activation. For each task represented in Fig 4C, cells within a condition were randomly split 80% / 10% / 10% into train / val / test splits, and probes were trained for five epochs before reporting final test classification accuracy.

To obtain AUROC and AUPR values, we used the State Embedding model to encode the Replogle-Nadig genetic perturbation screen dataset (Replogle et al., 2022; Nadig et al., 2025), then used the State decoder (Adduri et al., 2025) to reconstruct gene expression values from cell embeddings. For each perturbation, we computed effect scores as the absolute difference between mean predicted log probabilities (averaged across cells) under perturbed versus control conditions. Ground truth DEGs were identified using the Wilcoxon rank sum test at FDR < 0.05. ROC and PR curves were generated by varying the classification threshold across all unique effect score values in descending order, with genes above each threshold classified as differentially expressed and evaluated against the binary ground truth labels.

### 5.11. Cell type assignment

We retrieved and reprocessed Tabula Sapiens 10x Genomics data for 24 different tissues: blood, brain, bone marrow, lung, spleen, eye, liver, skin, muscle, small intestine, mammary, pancreas, lymph node, heart, large intestine, vasculature, fat, endometrium, prostate, bladder, trachea, salivary gland, tongue, and thymus. We also separately created a cell atlas for the brain based on a re-processing of the public data from the Human Brain cell atlas (Siletti et al., 2023). We used the State Embedding 600M (SE-600M) model to embed the Tabula Sapiens and Human Brain atlas data. For each tissue, we trained a separate logistic regression model to classify cell type based on the SE-600M embeddings. The logistic regression model was implemented using sklearn, with the max_iter parameter set to 1000.

To map samples (SRX, ERX) to the appropriate tissue model, we used the corresponding tissue category label from SRAgent. The raw text tissue labels of each sample were originally converted to one of 75 CELLx-GENE tissue categories by OpenAI’s o3 model, including ‘other’. We mapped each of these tissue categories in the metadata to the closest tissue model, and ‘other’ if it did not fit with any of the tissues. For each sample, we embedded the raw counts with the State Embedding model, and applied the classifier that was trained on the same tissue as the sample, making the assumption that each sample only contains one tissue. For our analysis, we omitted samples that had a mean UMI count per cell of 100 or less, and samples classified as ‘other’ tissue were not annotated. After these filters, we analyzed 166,381,968 human cells with 143 uniquely predicted cell types.

To visualize the cells across tissues, we randomly retrieved samples amounting to approximately 50,000 cells from each tissue, and we plotted a UMAP of these cells using the State embeddings. We colored one plot by the predicted cell type and one plot by the tissue label. Cells beyond the 0.1–99.9 percentile range for the x- and y-axis coordinates are excluded from the plot to filter out extreme outliers. We used the OBO Foundry Cell Ontology (Diehl et al., 2016) to color the UMAP with broader parent cell type categories that encompass our existing cell type labels.

To biologically validate our annotations, we compared cell type marker genes from PanglaoDB to the differentially expressed genes of our predicted cell types. To do this, we first mapped each of our cell type labels to a relevant cell type category in PanglaoDB. We randomly retrieved samples amounting to approximately 500,000 cells for each tissue in scBaseCount, or all of the cells for that tissue if there were less than 500,000 cells, and ranked the genes of each cell type category using t-test score as implemented in the scanpy `rank_genes_groups` function. We repeated this process for the cell type categories in each tissue of Tabula Sapiens. For each cell type category, we computed the percentage of PanglaoDB marker genes that were detected in the top 200 upregulated genes. We averaged this percentage for each cell type category across tissues, and we plotted these values from scBaseCount against Tabula Sapiens. The size of each point in the scatter plot reflects the number of markers in PanglaoDB for that cell type category. Each point is also colored by the Jaccard Similarity between the set of marker genes detected in scBaseCount and in Tabula Sapiens for that cell type category.

## 6. Acknowledgments

We especially thank Nianzhen Li for support and Jeremy Sullivan for managing infrastructure and compute resources. We thank Brian Plosky for important feedback in preparing the manuscript. We also thank Joseph Caputo, Julia Kazaks for their support. We would also like to thank Mingze Dong, Alishba Imran, Dhruv Gautam and Mohsen Naghipourfar for testing model training using scBaseCount. We thank the CZI CELLxGENE team for sharing SRA accession IDs to their datasets when available. We would like to thank Rahul Satija, Sourav Sarkar and Skylar Li for running early analysis on the scBaseCount data. S.K., P.D.H, and H.G. are Arc Core Investigators and acknowledge funding support from Arc Institute.

## 7. Author contributions

Y.R., A.D., and H.G. conceived the project. Y.R, H.G., A.D., P.D.H., S.K, and D.P.B. supervised the project. N.Y. conceived and developed `SRAgent`. N.Y., C.C., R.I, and A.D. developed `scRecounter`. C.C., N.Y., A.N., A.A., Y.R., J.P. and N.T. performed data analysis, incorporation of CZ CELLxGENE projects, and comparisons between the two datasets. N.Y., C.C., A.N., A.A. and C.R.-T. visualized data. A.A., R.S. trained AI models for learning representations of cells. Y.R., N.Y, C.C., A.A., A.N. and H.G. wrote the document, incorporating comments from all authors.

## 9. Supplementary figures

**Figure S1.**
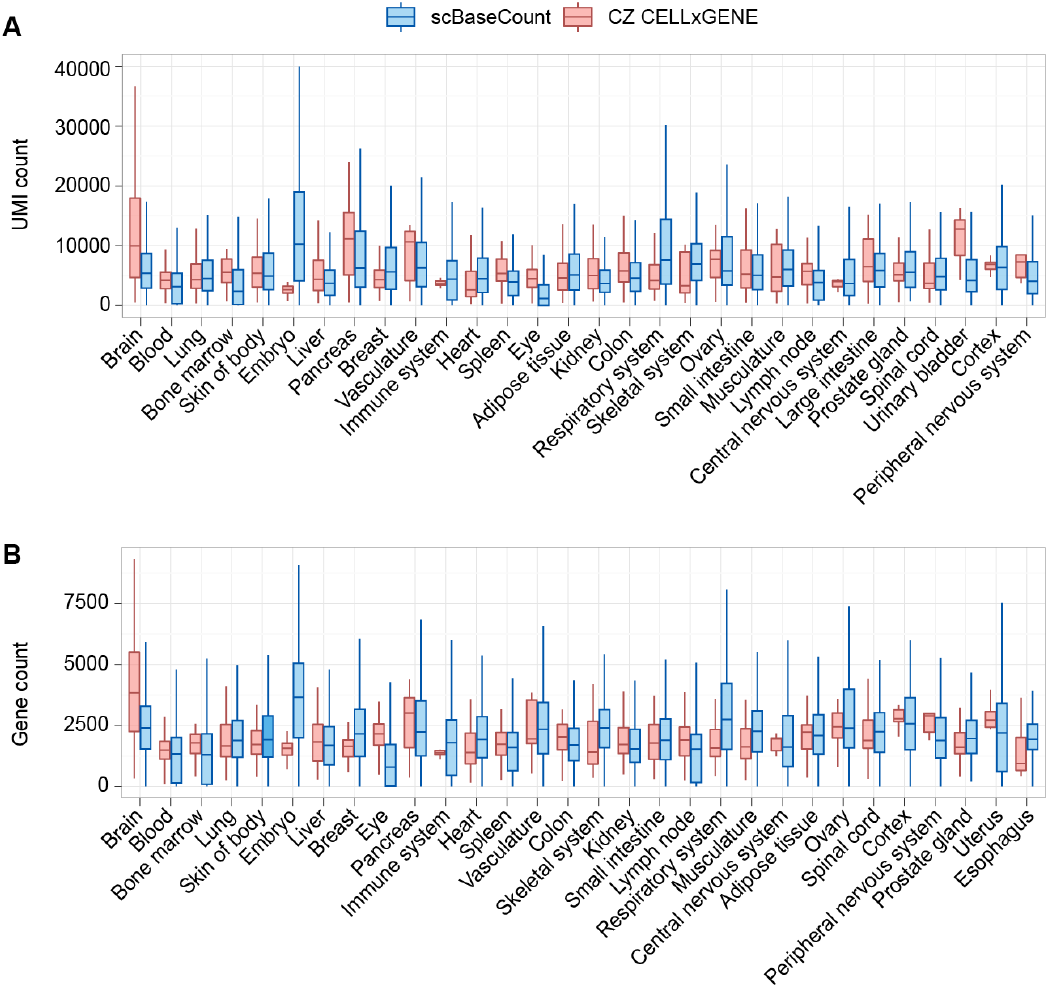
Count distribution comparison of scBaseCount to other publicly accessible single-cell data repositories. (**A-B**) Boxplots depicting the distribution of gene and UMI counts per cell across the top 30 tissues for scBaseCount (blue) and CZ CELLxGENE (red). The whiskers denote 1.5 × *IQR*. Just human and mouse cells were included.

**Figure S2.**
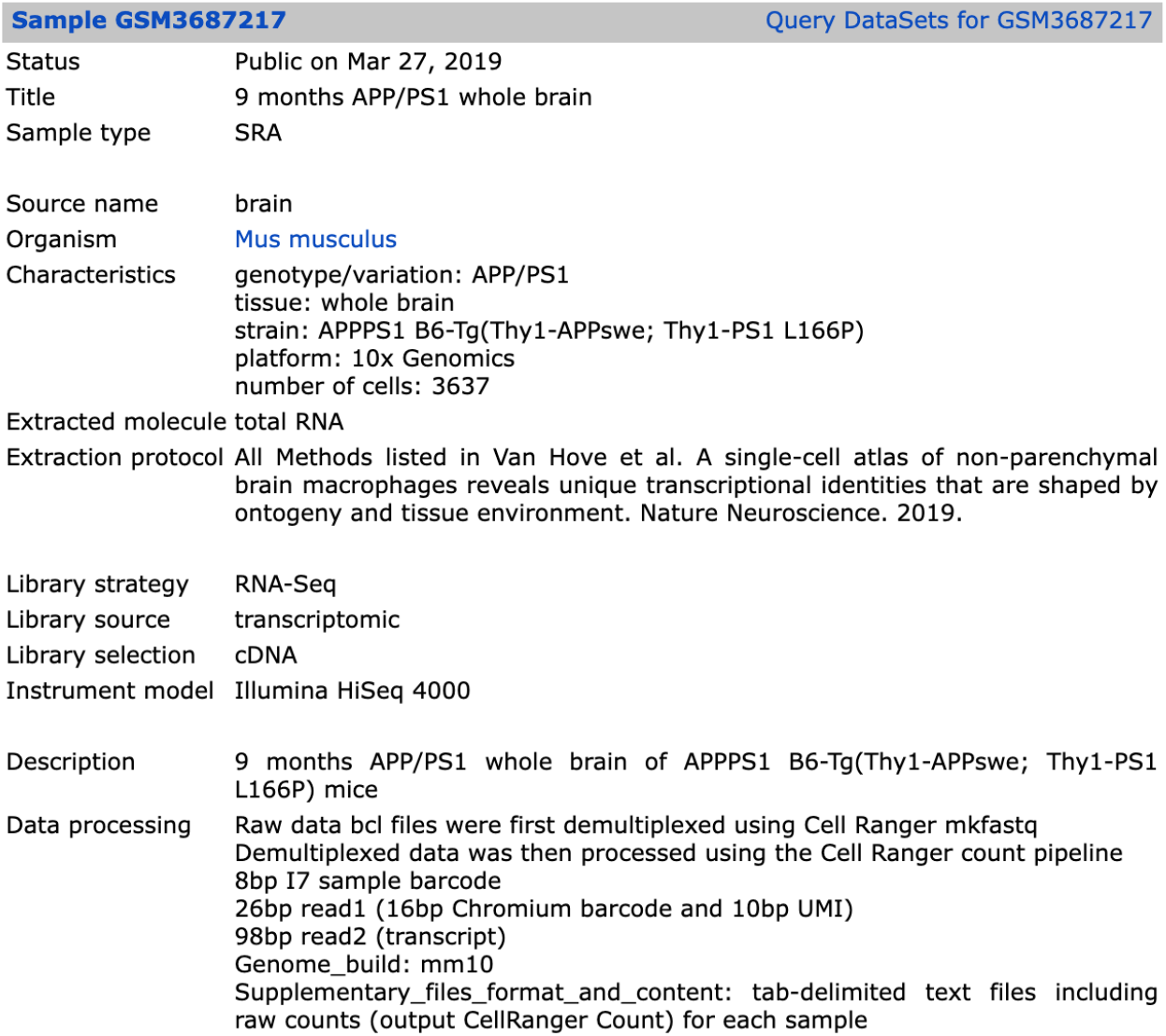
An example of an experiment where the disease label differs between scBaseCount and cellXgene. In scBaseCount, this sample is labeled as “Alzheimer’s”, due to the APP/PS1 mouse model, which is commonly used to study Alzheimer’s. In CELLxGENE, the cells from this experiment are labeled as healthy.

**Figure S3.**
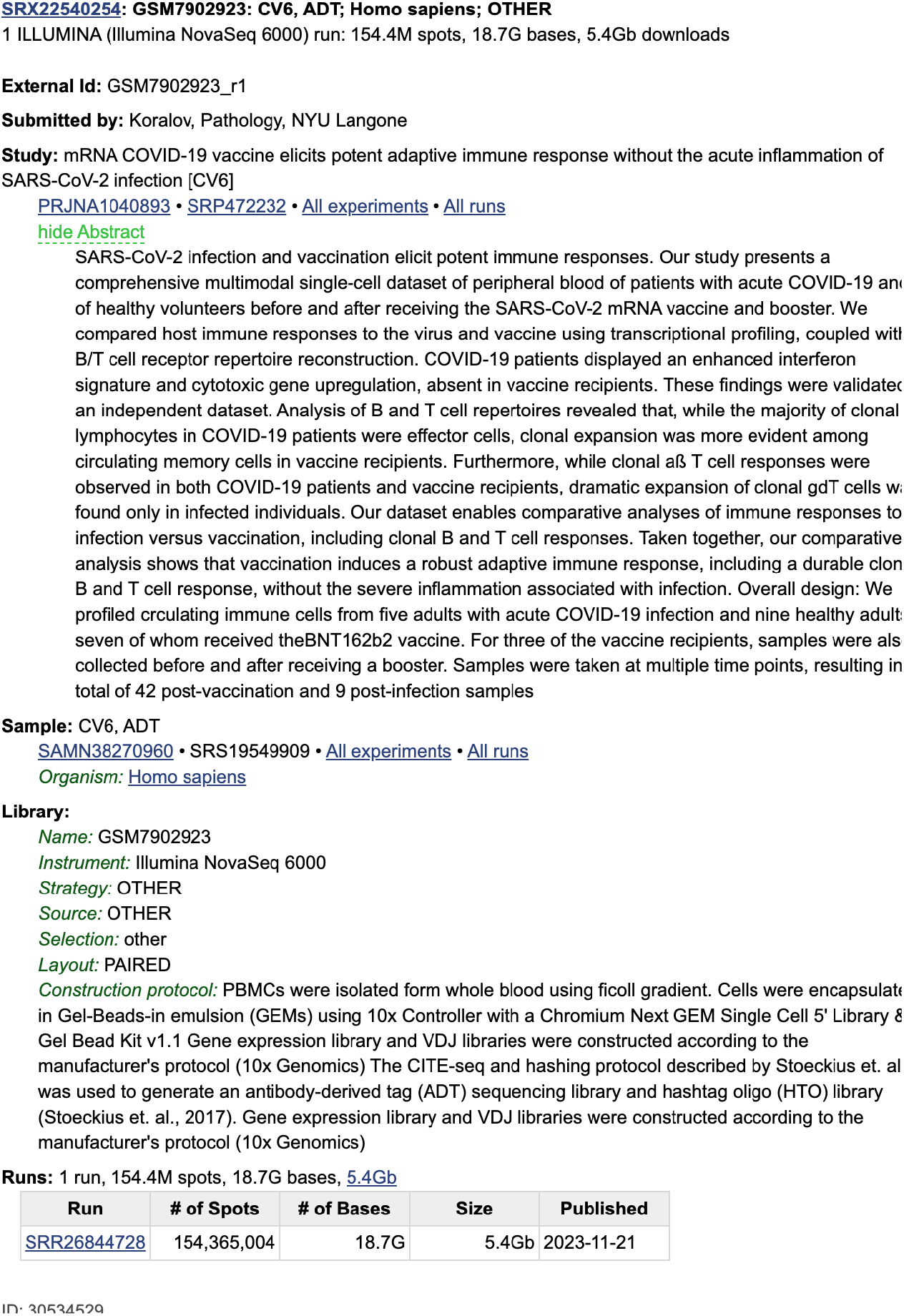
An example of an experiment where the disease label differs between scBaseCount and CELLxGENE. In scBaseCount, this sample is labeled as ‘Acute COVID’, whereas in CELLxGENE, the cells from this experiment are labeled as ‘normal’. This is assumed to be due to the fact that the control and treatment samples are not labeled in the metadata accessible to SRAgent.

**Figure S4.**
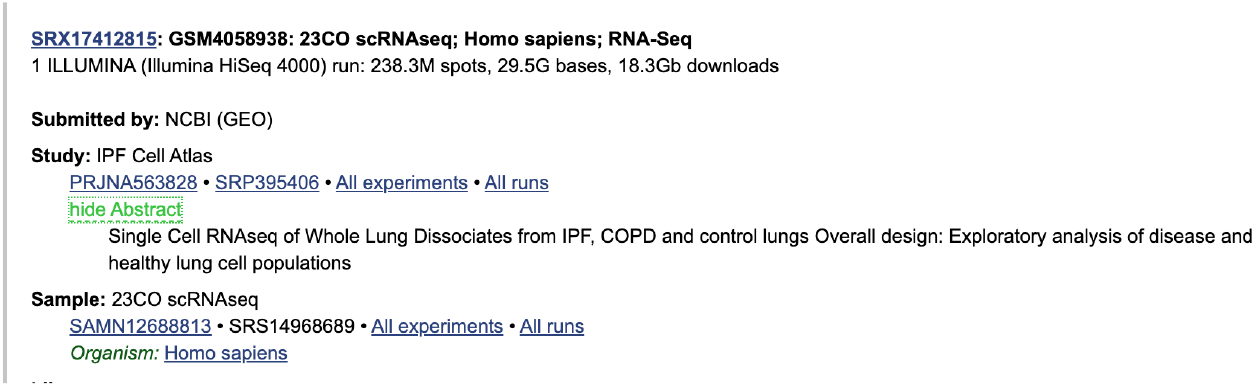
An example of an experiment where the disease label differs between scBaseCount and CELLxGENE. In scBaseCount, this sample is labeled as ‘COPD’, whereas in CELLxGENE, the cells from this experiment are labeled as ‘normal’. This is assumed to be due to the fact that the control and treatment samples are not labeled in the metadata accessible to SRAgent.

**Figure S5.**
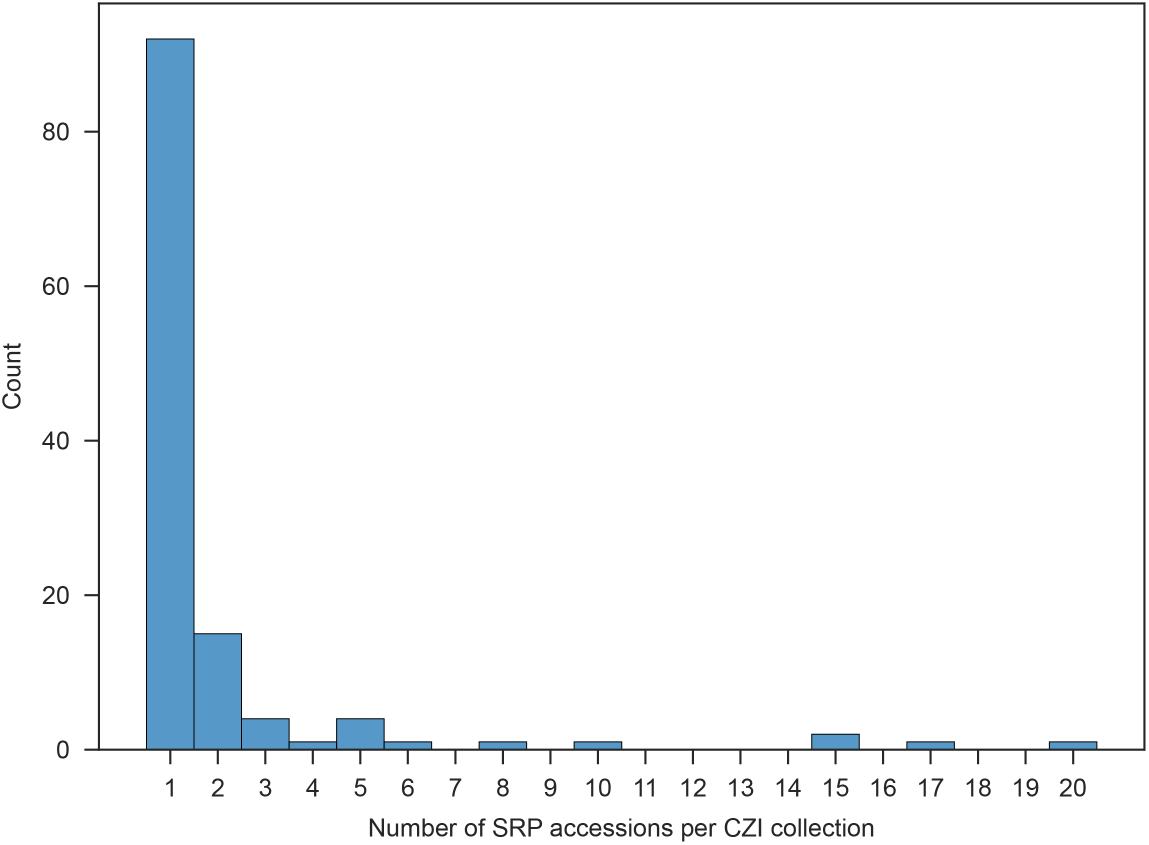
Histogram of the number of SRA studies (SRP) matching CZ CELLxGENE collection IDs. Since in most cases, there was a single SRP associated with a given CZI collection, we opted to treat these two variables as comparable in our data visualization in Figure 4.

**Figure S6.**
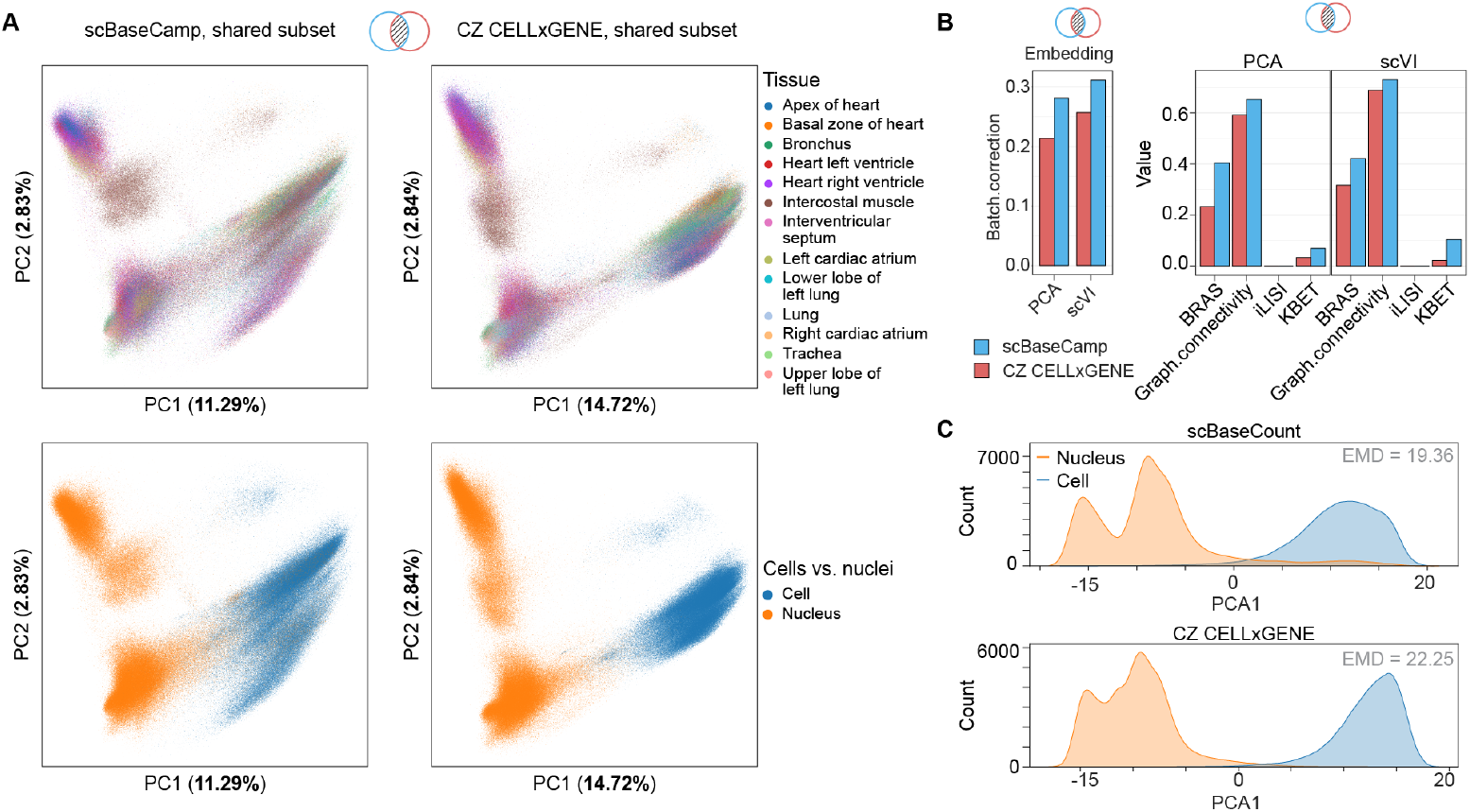
(**A**) Principal Component Analysis (PCA) of a shared subset of cells present in both scBaseCount and CZ CELLxGENE that contain both single cell and single nuclei data (N=566, 224). Left: PCA plots of the shared subset in scBaseCount, colored by tissue type (top) and suspension type (bottom). Right: PCA plots of the same subset in CZ CELLxGENE, showing a greater separation between single-cell and single-nucleus samples along the first two principal components. (**B**) SCIB batch correction metrics computed over PCA and scVI, for tissue type, and sample suspension type (i.e. cells vs nuclei), confirming that sample suspension type is a stronger driver of variation in CZ CELLxGENE than in scBaseCount, and tissue type is better preserved in scBaseCount. (**C**) Distribution of cells and nuclei projected onto the first PC for scBaseCount and CZ CELLx-GENE. We observed a more pronounced separation between cells and nuclei in CZ CELLxGENE as measured by Earth Mover’s Distance (EMD).

**Table S1.**
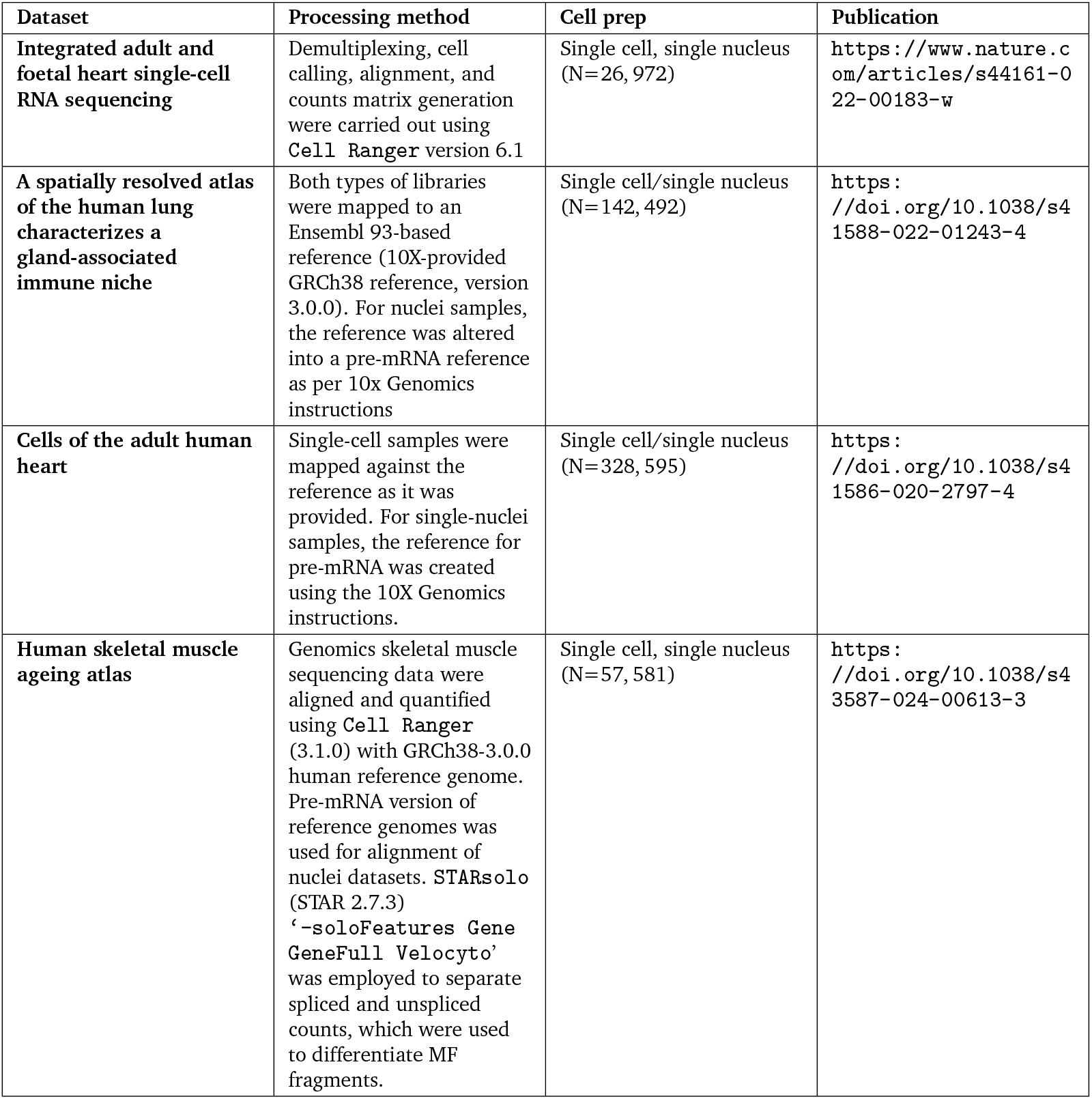
Overview of datasets, processing methods, and cell preparation techniques. Shared human single cell data collections in scBaseCount and CZ CELLxGENE containing both single-cell and single-nucleus sequencing data. The number of cells/nuclei in each dataset are shown in parentheses in the “Cell prep” column. As tabulated here, for single cell nuclei, the authors had used pre-mRNA references for read mapping, and default CellRanger mRNA mapping for single cell RNA-seq datasets. This difference in the choice of analytical approach contributes to the separation of single-cell and single-nuclei studies.

Table S2. **Overview of datasets, processing methods, and cell preparation techniques**. Shared human single cell data collections in scBaseCount and CZ CELLxGENE containing both single-cell and single-nucleus sequencing data. The number of cells/nuclei in each dataset are shown in parentheses in the “Cell prep” column. As tabulated here, for single cell nuclei, the authors had used pre-mRNA references for read mapping, and default CellRanger mRNA mapping for single cell RNA-seq datasets. This difference in the choice of analytical approach contributes to the separation of single-cell and single-nuclei studies.

**Table S3.**
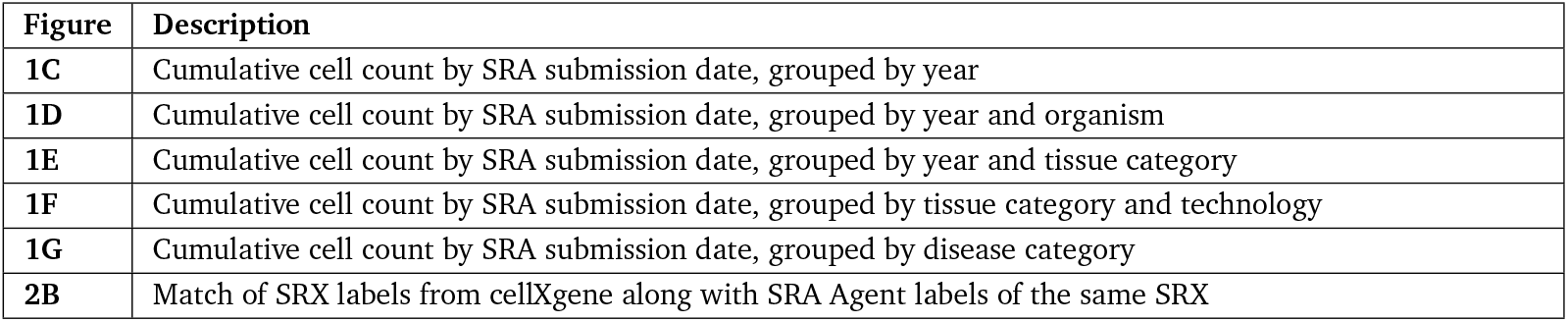
Overview of the supplemental tables. Each supplemental table provides the values used for plotting the figure listed in the ‘Figure’ column.

| Figure | Description |
| --- | --- |
| 1C | Cumulative cell count by SRA submission date, grouped by year |
| 1D | Cumulative cell count by SRA submission date, grouped by year and organism |
| 1E | Cumulative cell count by SRA submission date, grouped by year and tissue category |
| 1F | Cumulative cell count by SRA submission date, grouped by tissue category and technology |
| 1G | Cumulative cell count by SRA submission date, grouped by disease category |
| 2B | Match of SRX labels from cellXgene along with SRA Agent labels of the same SRX |

Table S4 | **Comparison of CZI and SRAgent annotations**. The attached CSV table presents the tissue and disease annotations from CZI (Chan Zuckerberg Initiative) alongside the corresponding SRAgent annotations for the same SRX accessions, highlighting annotation differences and standardization approaches between the two systems.

## References

A. K. Adduri, D. Gautam, B. Bevilacqua, A. Imran, R. Shah, M. Naghipourfar, N. Teyssier, R. Ilango, S. Nagaraj, M. Dong, C. Ricci-Tam, C. Carpenter, V. Subramanyam, A. Winters, S. Tirukkovular, J. Sullivan, B. S. Plosky, B. Eraslan, N. D. Youngblut, J. Leskovec, L. A. Gilbert, S. Konermann, P. D. Hsu, A. Dobin, D. P. Burke, H. Goodarzi, and Y. H. Roohani. Predicting cellular responses to perturbation across diverse contexts with state. bioRxiv, 2025. doi: 10.1101/2025.06.26.661135. URL https://www.biorxiv.org/content/early/2025/07/10/2025.06.26.661135.

C. Bunne, Y. Roohani, Y. Rosen, A. Gupta, X. Zhang, M. Roed, T. Alexandrov, M. AlQuraishi, P. Brennan, D.B. Burkhardt, et al. How to build the virtual cell with artificial intelligence: Priorities and opportunities. Cell, 187(25):7045–7063, 2024.

M. Büttner, Z. Miao, F. A. Wolf, S. A. Teichmann, and F. J. Theis. A test metric for assessing single-cell RNA-seq batch correction. Nat. Methods, 16(1):43–49, Jan. 2019.

Y. Chen and J. Zou. Simple and effective embedding model for single-cell biology built from ChatGPT. Nat. Biomed. Eng., pages 1–11, Dec. 2024.

T. T. S. Consortium*, R. C. Jones, J. Karkanias, M. A. Krasnow, A. O. Pisco, S. R. Quake, J. Salzman, N. Yosef, B. Bulthaup, P. Brown, et al. The tabula sapiens: A multiple-organ, single-cell transcriptomic atlas of humans. Science, 376(6594):eabl4896, 2022.

H. Cui, C. Wang, H. Maan, K. Pang, F. Luo, N. Duan, and B. Wang. scgpt: toward building a foundation model for single-cell multi-omics using generative ai. Nature Methods, 21(8):1470–1480, 2024.

CZI Cell Science Program, S. Abdulla, B. Aevermann, P. Assis, S. Badajoz, S. M. Bell, E. Bezzi, B. Cakir, J. Chaffer, S. Chambers, J. M. Cherry, T. Chi, J. Chien, L. Dorman, P. Garcia-Nieto, N. Gloria, M. Hastie, D. Hegeman, J. Hilton, T. Huang, A. Infeld, A.-M. Istrate, I. Jelic, K. Katsuya, Y. J. Kim, K. Liang, M. Lin, M. Lombardo, B. Marshall, B. Martin, F. McDade, C. Megill, N. Patel, A. Predeus, B. Raymor, B. Robatmili, D. Rogers, E. Rutherford, D. Sadgat, A. Shin, C. Small, T. Smith, P. Sridharan, A. Tarashansky, N. Tavares, H. Thomas, A. Tolopko, M. Urisko, J. Yan, G. Yeretssian, J. Zamanian, A. Mani, J. Cool, and A. Carr. CZ CELLxGENE discover: a single-cell data platform for scalable exploration, analysis and modeling of aggregated data. Nucleic Acids Res., 53(D1):D886–D900, Jan. 2025.

P. Di Tommaso, M. Chatzou, E. W. Floden, P. P. Barja, E. Palumbo, and C. Notredame. Nextflow enables reproducible computational workflows. Nat. Biotechnol., 35(4):316–319, Apr. 2017.

A. D. Diehl, T. F. Meehan, Y. M. Bradford, M. H. Brush, W. M. Dahdul, D. S. Dougall, Y. He, D. Osumi-Sutherland, A. Ruttenberg, S. Sarntivijai, C. E. Van Slyke, N. A. Vasilevsky, M. A. Haendel, J. A. Blake, and C. J. Mungall. The cell ontology 2016: enhanced content, modularization, and ontology interoperability. Journal of Biomedical Semantics, 7(1):44, Jul 2016. ISSN 2041-1480. doi: 10.1186/s13326-016-0088-7. URL https://doi.org/10.1186/s13326-016-0088-7.

A. Dobin, C. A. Davis, F. Schlesinger, J. Drenkow, C. Zaleski, S. Jha, P. Batut, M. Chaisson, and T. R. Gingeras. STAR: ultrafast universal RNA-seq aligner. Bioinformatics, 29(1):15–21, Jan. 2013.

C. Domínguez Conde, C. Xu, L. B. Jarvis, D. B. Rainbow, S. B. Wells, T. Gomes, S. Howlett, O. Suchanek, K. Polanski, H. King, et al. Cross-tissue immune cell analysis reveals tissue-specific features in humans. Science, 376(6594):eabl5197, 2022.

G. Eraslan, E. Drokhlyansky, S. Anand, E. Fiskin, A. Subramanian, M. Slyper, J. Wang, N. Van Wittenberghe, J. M. Rouhana, J. Waldman, O. Ashenberg, M. Lek, D. Dionne, T. S. Win, M. S. Cuoco, O. Kuksenko, A. M. Tsankov, P. A. Branton, J. L. Marshall, A. Greka, G. Getz, A.V. Segrè, F. Aguet, O. Rozenblatt-Rosen, K. G. Ardlie, and A. Regev. Single-nucleus cross-tissue molecular reference maps toward understanding disease gene function. Science, 376(6594):eabl4290, May 2022.

O. Franzén, L.-M. Gan, and J.L.M. Björkegren. Panglaodb: a web server for exploration of mouse and human single-cell rna sequencing data. Database, 2019:baz046, 04 2019. ISSN 1758-0463. doi: 10.1093/database/baz046. URL https://doi.org/10.1093/database/baz046.

A. C. Frazee, B. Langmead, and J. T. Leek. ReCount: a multi-experiment resource of analysis-ready RNA-seq gene count datasets. BMC Bioinformatics, 12(1):449, Nov. 2011.

S. Frölich, M. van der Sande, T. Schäfers, and S. J. van Heeringen. Genomepy: Genes and genomes at your fingertips. Bioinformatics, 39(3), Mar. 2023.

M. Hao, J. Gong, X. Zeng, C. Liu, Y. Guo, X. Cheng, T. Wang, J. Ma, X. Zhang, and L. Song. Large-scale foundation model on single-cell transcriptomics. Nature methods, 21(8):1481–1491, 2024.

G. Heimberg, T. Kuo, D. J. DePianto, O. Salem, T. Heigl, N. Diamant, G. Scalia, T. Biancalani, S. J. Turley, J. R. Rock, H. Corrada Bravo, J. Kaminker, J. A. Vander Heiden, and A. Regev. A cell atlas foundation model for scalable search of similar human cells. Nature, Nov. 2024.

W. Hou and Z. Ji. Assessing GPT-4 for cell type annotation in single-cell RNA-seq analysis. Nat. Methods, 21 (8):1462–1465, Aug. 2024.

A. Kazmi, D. Singh, S. Jatav, and S. Luthra. Beyond the hype: The complexity of automated cell type annotations with GPT-4. bioRxiv, page 2025.02.11.637659, Feb. 2025.

G. La Manno, R. Soldatov, A. Zeisel, E. Braun, H. Hochgerner, V. Petukhov, K. Lidschreiber, M. E. Kastriti, P. Lönnerberg, A. Furlan, et al. Rna velocity of single cells. Nature, 560(7719):494–498, 2018a.

G. La Manno, R. Soldatov, A. Zeisel, E. Braun, H. Hochgerner, V. Petukhov, K. Lidschreiber, M. E. Kastriti, P. Lönnerberg, A. Furlan, J. Fan, L. E. Borm, Z. Liu, D. van Bruggen, J. Guo, X. He, R. Barker, E. Sundström, G. Castelo-Branco, P. Cramer, I. Adameyko, S. Linnarsson, and P. V. Kharchenko. RNA velocity of single cells. Nature, 560(7719):494–498, Aug. 2018b.

M. Lange, V. Bergen, M. Klein, M. Setty, B. Reuter, M. Bakhti, H. Lickert, M. Ansari, J. Schniering, H. B. Schiller, D. Pe’er, and F. J. Theis. CellRank for directed single-cell fate mapping. Nat. Methods, 19(2):159–170, Feb. 2022.

R. Leinonen, H. Sugawara, M. Shumway, and International Nucleotide Sequence Database Collaboration. The sequence read archive. Nucleic Acids Res., 39(Database issue):D19–21, Jan. 2011.

B. Li, V. Ruotti, R. M. Stewart, J. A. Thomson, and C. N. Dewey. Rna-seq gene expression estimation with read mapping uncertainty. Bioinformatics, 26(4):493–500, 12 2009. ISSN 1367-4803. doi: 10.1093/bioinformatics/btp692. URL https://doi.org/10.1093/bioinformatics/btp692.

R. Lopez, J. Regier, M. B. Cole, M. I. Jordan, and N. Yosef. Deep generative modeling for single-cell transcriptomics. Nature methods, 15(12):1053–1058, 2018.

M. Lotfollahi, M. Naghipourfar, M. D. Luecken, M. Khajavi, M. Büttner, M. Wagenstetter, Ž. Avsec, A. Gayoso, N. Yosef, M. Interlandi, et al. Mapping single-cell data to reference atlases by transfer learning. Nature biotechnology, 40(1):121–130, 2022.

J. C. Melms, J. Biermann, H. Huang, Y. Wang, A. Nair, S. Tagore, I. Katsyv, A. F. Rendeiro, A. D. Amin, D. Schapiro, C. J. Frangieh, A. M. Luoma, A. Filliol, Y. Fang, H. Ravichandran, M. G. Clausi, G. A. Alba, M. Rogava, S. W. Chen, P. Ho, D. T. Montoro, A. E. Kornberg, A. S. Han, M. F. Bakhoum, N. Anandasabapathy, M. Suárez-Fariñas, S. F. Bakhoum, Y. Bram, A. Borczuk, X. V. Guo, J. H. Lefkowitch, C. Marboe, S. M. Lagana, A. Del Portillo, E. J. Tsai, E. Zorn, G. S. Markowitz, R. F. Schwabe, R. E. Schwartz, O. Elemento, A. Saqi, H. Hibshoosh, J. Que, and B. Izar. A molecular single-cell lung atlas of lethal COVID-19. Nature, 595(7865):114–119, July 2021.

C. J. Mungall, C. Torniai, G. V. Gkoutos, S. E. Lewis, and M. A. Haendel. Uberon, an integrative multi-species anatomy ontology. Genome Biology, 13(1):R5, 2012. doi: 10.1186/gb-2012-13-1-r5. URL https://genomebiology.biomedcentral.com/articles/10.1186/gb-2012-13-1-r5.

A. Nadig, J. M. Replogle, A. N. Pogson, M. Murthy, S. A. McCarroll, J. S. Weissman, E. B. Robinson, and L.J. O’Connor. Transcriptome-wide analysis of differential expression in perturbation atlases. Nature Genetics, pages 1–10, 2025.

J. D. Pearce, S. E. Simmonds, G. Mahmoudabadi, L. Krishnan, G. Palla, A.-M. Istrate, A. Tarashansky, B. Nelson, O. Valenzuela, D. Li, et al. A cross-species generative cell atlas across 1.5 billion years of evolution: The transcriptformer single-cell model. bioRxiv, pages 2025–04, 2025.

F. Pedregosa, G. Varoquaux, A. Gramfort, V. Michel, B. Thirion, O. Grisel, M. Blondel, P. Prettenhofer, R. Weiss, V. Dubourg, J. Vanderplas, A. Passos, D. Cournapeau, M. Brucher, M. Perrot, and E. Duchesnay. Scikit-learn: Machine learning in Python. Journal of Machine Learning Research, 12:2825–2830, 2011.

N. G. Ravindra, M. M. Alfajaro, V. Gasque, N. C. Huston, H. Wan, K. Szigeti-Buck, Y. Yasumoto, A. M. Greaney, V. Habet, R. D. Chow, et al. Single-cell longitudinal analysis of sars-cov-2 infection in human airway epithelium identifies target cells, alterations in gene expression, and cell state changes. PLoS biology, 19(3): e3001143, 2021.

J. M. Replogle, R. A. Saunders, A. N. Pogson, J. A. Hussmann, A. Lenail, A. Guna, L. Mascibroda, E. J. Wagner, K. Adelman, G. Lithwick-Yanai, et al. Mapping information-rich genotype-phenotype landscapes with genome-scale perturb-seq. Cell, 185(14):2559–2575, 2022.

J. E. Rood, A. Maartens, A. Hupalowska, S. A. Teichmann, and A. Regev. Impact of the human cell atlas on medicine. Nat. Med., 28(12):2486–2496, Dec. 2022.

Y. H. Roohani, T. J. Hua, P.-Y. Tung, L. R. Bounds, F. B. Yu, A. Dobin, N. Teyssier, A. Adduri, A. Woodrow, B. S. Plosky, et al. Virtual cell challenge: Toward a turing test for the virtual cell. Cell, 188(13):3370–3374, 2025.

Y. Rosen, Y. Roohani, A. Agarwal, L. Samotorčan, T. S. Consortium, S. R. Quake, and J. Leskovec. Universal cell embeddings: A foundation model for cell biology. bioRxiv, pages 2023–11, 2023.

O. Rozenblatt-Rosen, M. J. T. Stubbington, A. Regev, and S. A. Teichmann. The human cell atlas: from vision to reality. Nature, 550(7677):451–453, Oct. 2017.

K. Siletti, R. Hodge, A. M. Albiach, K. W. Lee, S.-L. Ding, L. Hu, P. Lönnerberg, T. Bakken, T. Casper, M. Clark, N. Dee, J. Gloe, D. Hirschstein, N. V. Shapovalova, C. D. Keene, J. Nyhus, H. Tung, A. M. Yanny, E. Arenas, E. S. Lein, and S. Linnarsson. Transcriptomic diversity of cell types across the adult human brain. Science, 382(6667):eadd7046, 2023. doi: 10.1126/science.add7046. URL https://www.science.org/doi/abs/10.1126/science.add7046.

C. V. Theodoris, L. Xiao, A. Chopra, M. D. Chaffin, Z. R. Al Sayed, M. C. Hill, H. Mantineo, E. M. Brydon, Z. Zeng, X. S. Liu, et al. Transfer learning enables predictions in network biology. Nature, 618(7965):616–624, 2023.

M. G. van der Wijst, S. E. Vazquez, G. C. Hartoularos, P. Bastard, T. Grant, R. Bueno, D. S. Lee, J. R. Greenland, Y. Sun, R. Perez, et al. Longitudinal single-cell epitope and rna-sequencing reveals the immunological impact of type 1 interferon autoantibodies in critical covid-19: Anti-ifn antibodies in critical covid-19 correlate with poor isg response and upregulation of lair1 surface protein in pbmcs. BioRxiv, 2021.

N. A. Vasilevsky, N. A. Matentzoglu, S. Toro, J. E. F. IV, H. B. Hegde, D. R. Unni, G. F. Alyea, J. S. Amberger, L. Babb, J. P. Balhoff, T. I. Bingaman, G. A. Burns, T. J. Callahan, L. C. Carmody, L. E. Chan, G. S. Chang, M. Dumontier, L. E. Failla, M. J. Flowers, H. A. Garrett, D. Gration, T. Groza, M. Hanauer, N. L. Harris, I. Helbig, J. A. Hilton, D. S. Himmelstein, C. T. Hoyt, M. S. Kane, S. Köhler, D. Lagorce, M. Larralde, A. Lock, I. L. Santiago, D. R. Maglott, A. J. Malheiro, B. H. M. Meldal, J. A. McMurry, M. Munoz-Torres, T. H. Nelson, D. Ochoa, T. I. Oprea, D. Osumi-Sutherland, H. Parkinson, Z. M. Pendlington, A. Rath, H. L. Rehm, L. Remennik, E. R. Riggs, P. Roncaglia, J. E. Ross, M. F. Shadbolt, K. A. Shefchek, M. N. Similuk, N. Sioutos, R. Sparks, R. Stefancsik, R. Stephan, D. Stupp, J. C. Sundaramurthi, I. Tammen, D. Tay, C. L. Thaxton, E. Valasek, J. Valls-Margarit, A. H. Wagner, N. L. Washington, C. Bizon, M. A. Haendel, P. N. Robinson, and C. J. Mungall. Mondo: Unifying diseases for the world, by the world. medRxiv, 2022. doi: 10.1101/2022.04.13.22273750. URL https://www.medrxiv.org/content/10.1101/2022.04.13.22273750v3. Preprint.

T. T.-H. Wu, K. J. Travaglini, A. Rustagi, D. Xu, Y. Zhang, L. Andronov, S. Jang, A. Gillich, R. Dehghannasiri, G.J. Martínez-Colón, et al. Interstitial macrophages are a focus of viral takeover and inflammation in covid-19 initiation in human lung. Journal of Experimental Medicine, 221(6):e20232192, 2024.

Y. Xiao, J. Liu, Y. Zheng, X. Xie, J. Hao, M. Li, R. Wang, F. Ni, Y. Li, J. Luo, S. Jiao, and J. Peng. Cel-lAgent: An LLM-driven multi-agent framework for automated single-cell data analysis. bioRxiv, page 2024.05.13.593861, May 2024.

S. Yao, J. Zhao, D. Yu, N. Du, I. Shafran, K. Narasimhan, and Y. Cao. ReAct: Synergizing reasoning and acting in language models. arXiv [cs.CL], Oct. 2022.

